# CD47 prevents the elimination of diseased fibroblasts in scleroderma

**DOI:** 10.1101/2020.06.06.138222

**Authors:** Tristan Lerbs, Lu Cui, Megan E. King, Tim Chai, Claire Muscat, Tyler Shibata, Gerlinde Wernig

## Abstract

Scleroderma is a devastating fibrotic autoimmune disease. Current treatments are partly effective in preventing disease progression, but do not remove fibrotic tissue. Here, we evaluated whether scleroderma fibroblasts take advantage of the “don’t-eat-me-signal” CD47 and whether blocking CD47 enables the body’s immune system to get rid of diseased fibroblasts. To test this approach, we used a *Jun*-inducible scleroderma model. We first demonstrated in patient samples that scleroderma upregulated JUN and increased promotor accessibilities of both JUN and the CD47. Next, we established our scleroderma model demonstrating that *Jun* mediated skin fibrosis through the hedgehog-dependent expansion of CD26+Sca1-fibroblasts in mice. In a niche-independent adaptive transfer model, JUN steered graft survival and conferred increased self-renewal to fibroblasts. In vivo, JUN enhanced the expression of CD47, and inhibiting CD47 eliminated an ectopic fibroblast graft and increased in vitro phagocytosis. In the syngeneic mouse, depleting macrophages ameliorated skin fibrosis. Therapeutically, combined CD47 and IL6 blockade reversed skin fibrosis in mice and led to the rapid elimination of ectopically transplanted scleroderma cells. Altogether, our study is the first to demonstrate the efficiency of combining different immunotherapies in treating scleroderma and provide a rationale for combining CD47 and IL6 inhibition in clinical trials.

## Introduction

Scleroderma is the fibrotic skin manifestation of the autoimmune diseases morphea and systemic sclerosis, causing significant morbidity. Current treatments rely on immune suppression and primarily aim at stopping or slowing down the progression of scleroderma [1, 2]. Unfortunately, these treatments only bring temporary relief to most patients. Therefore, we aimed at exploring a novel therapeutic approach that could not only prevent but also reverse the fibrotic skin changes. Considering that scleroderma leads to the accumulation of diseased fibroblasts, we hypothesized that the inhibition of the “don’t-eat-me-signal” CD47 would help the body’s immune system to remove abnormal fibroblasts. This therapeutic approach would not only stop the disease as immune suppression does but could permit its remission, thereby offering a breakthrough for the treatment of scleroderma.

Next to changes in the immune and vascular system, skin fibrosis represents one of scleroderma’s clinical symptoms [3–5]. Fibrosis is marked by an excessive amount of connective tissue primarily formed by fibroblasts. One of the genes that fibroblasts commonly upregulate in several fibrotic conditions, especially idiopathic pulmonary fibrosis, is the transcription factor *JUN* as our group has recently shown [6, 7]. *JUN*, that belongs to the family of the AP-1 transcription factors and that is activated through phosphorylation (pJUN), otherwise contributes to malignant diseases and plays a role in different developmental programs [8, 9]. To study the effects of *Jun*, our group uses a unique *Jun*-driven mouse model. In this model, *Jun* is inserted into the collagen locus and is under the control of a tetracycline-dependent promoter in the Rosa26 locus. Through doxycycline application, *Jun* can be induced either systemically through intraperitoneal injections or the drinking water, or locally through injections. Another characteristic feature of fibroblasts is their phenotypic and functional diversity [10]. During development, they form different lineages. Lineages can be distinguished by specific surface markers and both CD26 and Sca1 are among these markers. While CD26+/Sca1-fibroblasts primarily reside in the upper dermis, CD26-/Sca1+ fibroblasts can be mainly found in the lower dermis [10]. Though not specifically explored in these subsets of fibroblasts, hedgehog signaling with its main effector *Gli1* has been shown to contribute to dermal fibrosis [11]. Besides, hedgehog signaling plays a vital role in development and the maintenance of stem cell population [12, 13].

Immune therapy has changed the therapeutic landscapes for several cancers within a few years [14], unleashing the body’s own immune system in its fight against the tumor. PD1/PDL1 inhibition boosts T cell function and is approved as a first-line therapy for various cancers including NSCLC [15, 16]. CD47 prevents macrophages from eating their target cell and CD47 inhibition is currently investigated in advanced clinical trials [17, 18]. In contrast to immune checkpoint inhibitors and “don’t-eat-me-signals”, interleukin 6 blockade aims at reducing inflammatory responses that can worsen autoimmune diseases [19].

Conceptually, this study aimed at investigating whether the “don’t-eat-me-signal” CD47 either alone or in combination with interleukin 6 blockade is able to prevent or reverse fibrotic skin changes in a *Jun*-driven mouse model. We hypothesized that this combination would the immune system allow to get rid of abnormal fibroblasts while preventing an ongoing inflammatory response through interleukin 6 blockade.

## Results

### Human scleroderma activates JUN and CD47

At the beginning, we determined if JUN, CD47 and PDL1 are commonly upregulated in human scleroderma. Staining tissue sections, almost all FSP1+ fibroblasts in scleroderma but only a minority of FSP1+ fibroblasts in normal skin expressed pJUN (Fig. 1A, B). We then evaluated how primary dermal fibroblasts from scleroderma and normal skin regulate promotor accessibility of *JUN* and the hedgehog-associated genes *GLI1* and *PTCH1* through ATAC Seq studies (Fig. 1 C). In addition, we ran these studies after *JUN* knockout and under vismodegib to study the effects of *JUN* and hedgehog inhibition in scleroderma fibroblasts (Fig. 1 C). In accordance with our hypothesis, promotor accessibilities of *JUN*, *GLI1* and *PTCH* were increased in scleroderma compared to normal skin (Fig. 1 C). Vice versa, knocking *JUN* out decreased the promotor accessibility of *GLI1* and *PTCH1*. As the focus of our study was to evaluate immune therapy, we analyzed the ATAC Seq data for *CD47*, *PDL1* and *IL6* as well (Fig. 1 D). Promotor accessibility of all three genes was increased in scleroderma fibroblasts and *JUN* knockout led to decreased promotor accessibilities of *CD47*, *PDL1* and *IL6*. Finally, ATAC Seq demonstrated distinct promotor accessibility clustering before and after *JUN* knockout in scleroderma fibroblasts and increased promotor accessibilities of several fibrosis-associated genes (Fig. 1 E, F). Immunostainings of human scleroderma samples confirmed the expression of CD47, CD26 and PD1 (Fig. S1). The ATAC Seq studies did not answer which mechanism leads to the activation of JUN in scleroderma fibroblasts. One characteristic of the scleroderma skin, that it shares with all other fibrotic diseases, is its increased stiffness [20]. To test if stiffer conditions themselves induce JUN activation, we plated primary scleroderma fibroblasts either on a stiff 70kDa hydrogel or on a regular polystyrene dish. After two days, pJUN was significantly increased on the hydrogel, a result that we confirmed with primary pulmonary fibroblasts (Fig. 1 G). In conclusion, our results with patient sections and human fibroblasts indicated that JUN was activated in human scleroderma both on the protein and the molecular level, that JUN interacts with the hedgehog pathway, and stiffer conditions on a hydrogel themselves induce JUN activation.

**Figure 1.**
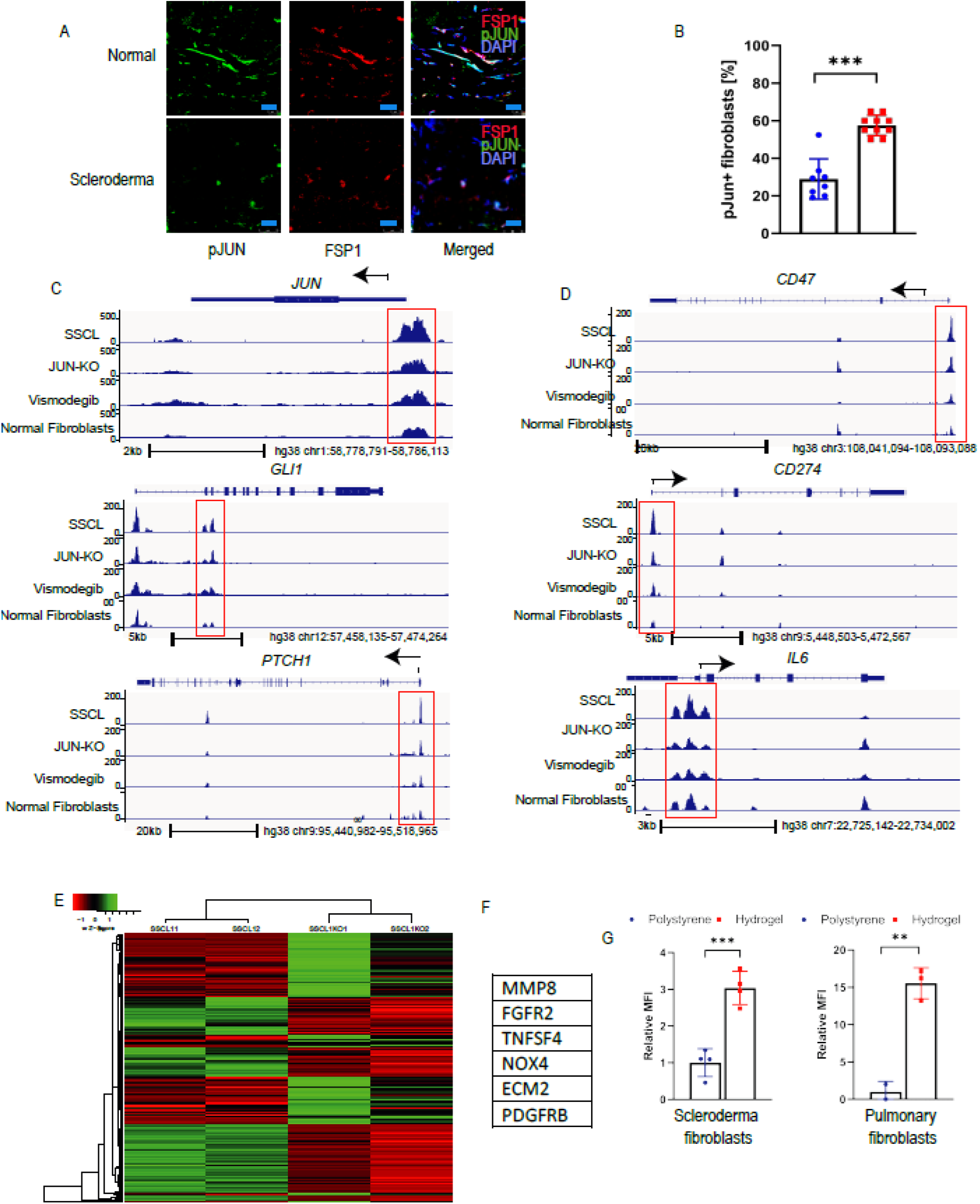
Human scleroderma activates JUN and CD47. **(A)** IF pictures of human scleroderma and normal skin stained against pJUN and FSP1, counterstained with DAPI. Scale bar = 25 µm. n=8-10. **(B)** Corresponding quantification of pJUN+ fibroblasts. Two-sided t-test. *** p < 0.001. n=8-10. Bar graphs represent means with standard deviations. **(C)** ATAC Seq analysis for JUN and the hedgehog genes Gli1 and Ptch1 in scleroderma fibroblasts (SSCL), scleroderma fibroblasts after JUN knockout (JUN-KO) or under vismodegib, or normal skin fibroblasts. The promoter regions are highlighted with red boxes. n=2. **(D)** ATAC Seq analysis for the immune checkpoints CD47 and PDL1 and the interleukin IL6 in scleroderma fibroblasts (SSCL), scleroderma fibroblasts after JUN knockout (JUN-KO) or under vismodegib, or normal skin fibroblasts. The promoter regions are highlighted with red boxes. n=2. **(E)** Heatmap of differential open chromatin regulatory elements characterized from ATAC-seq. The color bar shows the relative ATAC-seq signal (Z score of normalized read counts) as indicated. Samples 1 and 2 in both groups are individual samples. n=2. **(F)** Fibrosis-linked genes with a fivefold decline in promoter accessibility after JUN knockout. n=2.| **(G)** pJUN expression in pulmonary fibroblasts and scleroderma fibroblasts on a 70 kPa hydrogel or a regular polystyrene plastic dish. Two-sided t-test. ** p < 0.01 *** p < 0.001. n=4. Turkey’s multiple comparisons test. n=4. Bar graphs represent means with standard deviations.

### Jun expands distinct fibroblast populations in a scleroderma model

Before studying the efficiency of CD47, we characterized the mechanisms of our *Jun*-driven mouse model, and how it compared to the widely used bleomycin model (Fig. S2 A) [21]. We either induced *Jun* through intradermal doxycycline administrations every other day or administered intradermal bleomycin once at the beginning. After two weeks, both treatments led to significant fibrotic changes and a loss of fat tissue in the skin (Fig. 2 A, B). Fibrotic changes included the whole dermis and the tissue below the subcutaneous muscle tissue (Fig. S2 B-D). Interestingly, both *Jun* induction and bleomycin administration induced JUN activation to the same extent (Fig. 2 B, S3 A, B). JUN additionally led to an increased inflammatory infiltrate (Fig. S3) We then evaluated how *Jun* and bleomycin affect distinct fibroblast populations. For this purpose, we divided CD45-CD31-CD326-dermal fibroblasts into four groups, dependent on their expression of CD26 and SCA1 (Fig. 2 C, S4). Both *Jun* induction and bleomycin expanded CD26+/SCA1-fibroblasts (CD26+ fibroblasts) and decreased CD26+/Sca1+ (double-positive, DP) fibroblasts (Fig. 2 D, E). In a time-course study over five days, *Jun* increased CD26+ fibroblasts gradually while it decreased DP fibroblasts directly at the beginning (Fig. S5 A - C). When *Jun* is induced longer than five days, CD26+ fibroblasts started to decrease while CD26-/SCA1-(double-negative, DN) fibroblasts expanded (Fig. S5 D). As proliferation did not specifically affect CD26+ and CD26-fibroblasts this did suggest that immature CD26+ fibroblasts differentiate into mature DN fibroblasts (Fig. S5 E, F).

**Figure 2.**
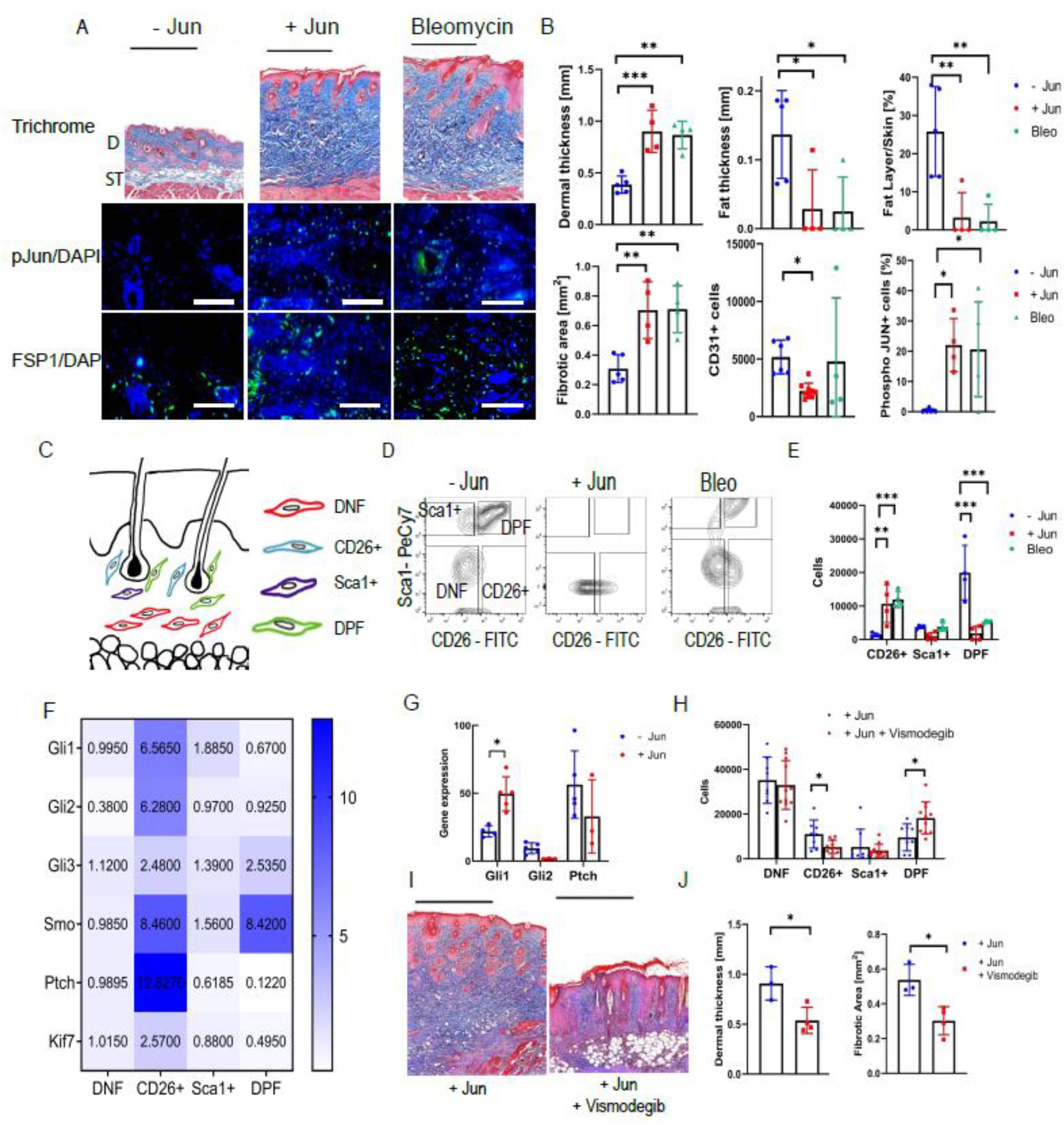
JUN expands distinct fibroblast populations in a hedgehog dependent manner. **(A)** Representative trichrome stains and immunofluorescence stains against pJUN and FSP1 without JUN induction (-JUN), with JUN induction (+JUN) and after bleomycin injection. Black scale bar = 500 µm. White scale bar = 75 µm. **(B)** Corresponding quantification of dermal thickness, fat layer thickness, fat/skin thickness relationship, fibrotic areas, total CD31+ endothelial cells and dermal pJUN+ cells/Field of view. Turkey’s multiple comparisons test. * p < 0.05 ** p < 0.01 *** p < 0.001. n=4-8. Bar graphs represent means with standard deviations. **(C)** Schematic drawing of the localization of double-positive fibroblasts (DPF), CD26+ fibroblasts, Sca1+ fibroblasts and double-negative fibroblasts (DNF) **(D)** Representative fibroblast FACS plots without JUN induction, with JUN induction and after bleomycin injection **(E)** Corresponding quantification of total CD26+ fibroblasts, Sca1+ fibroblasts and DPF. Turkey’s multiple comparisons test. *** p < 0.001. n=4. Bar graphs represent means with standard deviations. **(F)** Heatmap of the expression of different hedgehog-associated genes in the different fibroblast populations. Values are normalized to the expression in DNF. n=3-6. **(G)** Expression of Gli1, Gl2 and Ptch after JUN induction. All values are compared to the same standard value. Two-sided t-test. * p < 0.05. n=3-6. Bar graphs represent means with standard deviations. **(H)** Fibroblast populations after three days of Jun induction +/- hedgehog inhibition. Indicated are total cells/100,000 live cells. Fisher’s multiple comparisons test. * p < 0.05 ** p < 0.01. n=8-11. Bar graphs represent means with standard deviations. **(I)** Representative trichrome stainings after intradermal JUN induction +/- hedgehog inhibition. Scale = 500 µm. **(J)** Corresponding quantification of dermal thickness and fibrotic area. Two-sided t-test. * p<0.05. n=3. Bar graphs represent means with standard deviations.

### CD26+ fibroblast expansion and skin fibrosis is hedgehog-dependent

After demonstrating that JUN causes dermal fibrosis and expands distinct CD26+ fibroblasts, we evaluated whether JUN distinctively influences hedgehog signaling in dermal fibroblast populations. Comparing the expression of several hedgehog-associated genes in FACS purified dermal fibroblasts, we observed increased hedgehog activation in CD26+ fibroblasts and inducing *Jun* even further increased the expression of the main hedgehog effector *Gli*1 (Fig. 2 F, G, S6). To test if the hedgehog activation is mandatory for the expansion of CD26+ fibroblasts and the fibrotic skin changes under JUN, we blocked the hedgehog pathway with the smoothened inhibitor vismodegib [22]. Hedgehog inhibition did not only reduce the expansion of CD26+ fibroblasts but also almost completely prevented skin fibrosis after two weeks (Fig. 2 H, I, J). In conclusion, these results suggested that *Jun* drives skin fibrosis through the distinct activation of hedgehog signaling in CD26+ fibroblasts.

### Jun mediates increased self-renewal to fibroblasts

As a last step before evaluating CD47, we evaluated the effect of *Jun* on dermal fibroblasts outside their dermal niche. For this purpose, we first isolated fibroblasts from neonatal mouse skin. In vitro, an EdU uptake and subsequent flow cytometry demonstrated increased cell proliferation under *Jun* induction and in accordance with increased cell proliferation, JUN increased pStat3 signaling (Fig. 3 A, B, C). To investigate how JUN effects cell survival and proliferation in an niche-independent in vivo environment, we used an adaptive transfer model in which we transplanted GFP/Luciferase-labeled primary mouse dermal fibroblasts under the kidney capsule of immunocompromised NOD.Scid.Gamma (NSG) mice [23]. Tracking transplanted cells through luciferase-based optical imaging, knocking *Jun* out decreased graft survival (Fig. 3 D). When comparing non-*Jun* induced cells to *Jun*-induced cells, *Jun* increased cell proliferation both in optical imaging and histology (Fig. 3 E, F, G). Based on these and our previous results, we hypothesized that *Jun* increases self-renewal in fibroblasts. Supporting this hypothesis, blocking the stem cell-associated hedgehog pathway eliminated the fibroblast graft (Fig. 3 H, I). We additionally explored a serial transplantation model in which we transplanted RFP-labeled fibroblasts directly from one mouse to the next (Fig. 3 J). While we could no longer detect any RFP-labeled cells after the second recipient without *Jun* induction, we observed RFP-labeled cells even in the fourth recipient with *Jun* induction (Fig. 3 K). This result strongly supports that *Jun* mediates increased self-renewal to fibroblasts.

**Figure 3.**
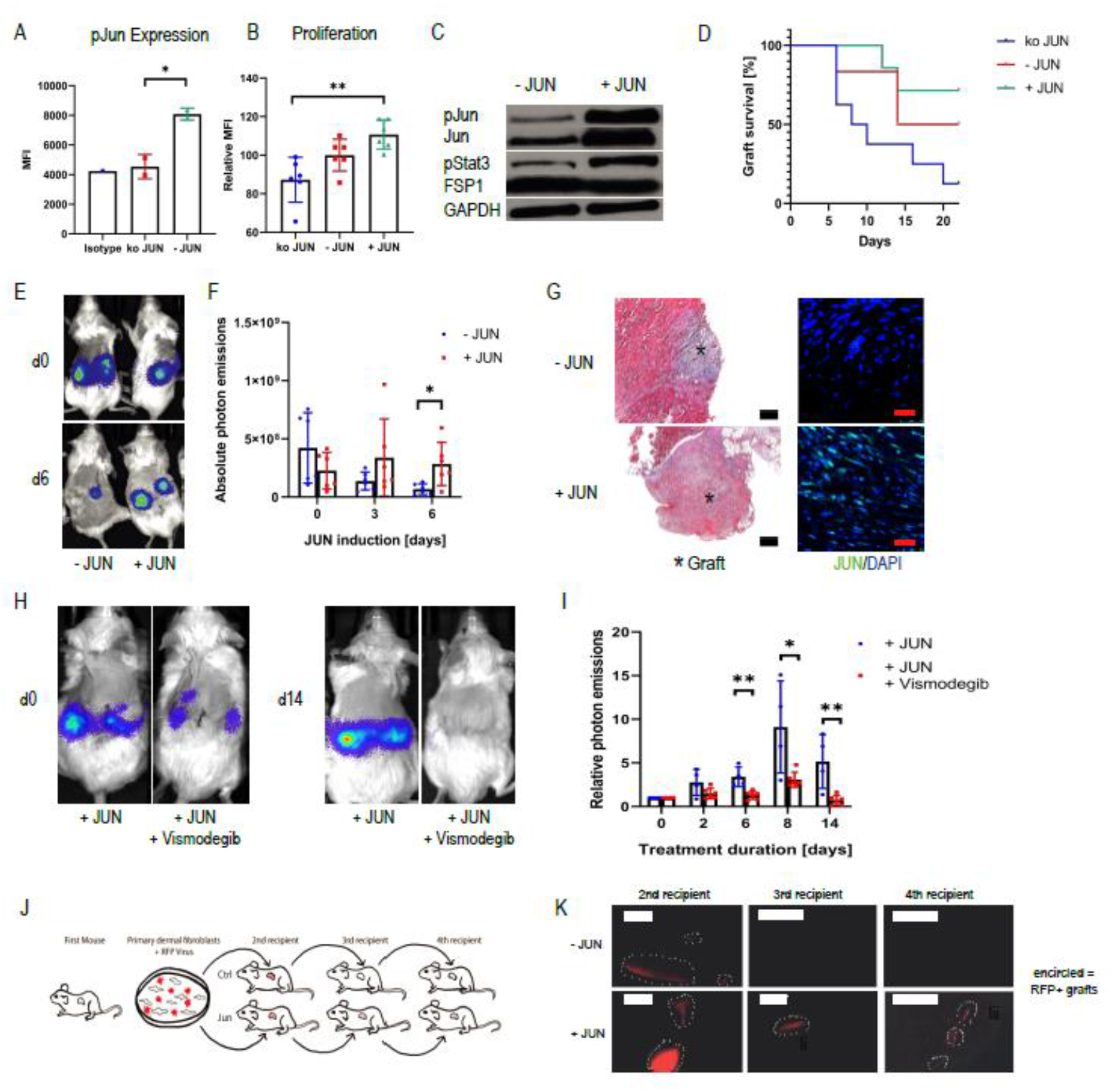
JUN mediates increased self-renewal to fibroblasts. **(A)** Mean Fluorescence Intensity for pJUN after JUN knockout (ko JUN). Turkey’s multiple comparisons test * p < 0.05. n=2. Bar graphs represent means with standard deviations. **(B)** Mean fluorescence intensity for incorporated EdU (AF594) after JUN knockout (ko JUN), without JUN induction (- JUN) and under JUN induction (+ JUN). Turkey’s multiple comparisons test. ** p < 0.01. n=6. Bar graphs represent means with standard deviations. **(C)** Western blot bands for pJUN, pStat3, FSP1 and GAPDH without and with JUN induction in primary mouse dermal fibroblasts **(D)** Kaplan Meier Curve of adaptive transfer graft survivals with JUN induction (+JUN), without JUN induction (- JUN) and after JUN knockout (ko JUN). Photon emissions below 100,000 were considered as a lost graft. n=5-8. **(E)** Representative optical images of an adaptive transfer model with JUN inducible fibroblasts. n=4-6. **(F)** Quantification of absolute photon emissions with and without JUN induction. Fisher’s multiple comparisons test. * p < 0.01. n=4-6. Bar graphs represent means with standard deviations. **(G)** Corresponding trichrome (10x) and pJUN stains of grafts. Black scale bar = 200 µm. Red scale bar = 25 µm. n=4-6. **(H)** Representative optical images of a JUN inducible adaptive transfer model +/- vismodegib. n=4-6. **(I)** Corresponding quantification of photon emissions. Values were normalized to the expression at day 0. Fisher’s multiple comparisons test. * p < 0.05 ** p < 0.01. n=4-6. Bar graphs represent means with standard deviations. **(J)** Schema of the adaptive serial transplantation model **(K)** Corresponding pictures of RFP+ cells in the 2^nd^, 3^rd^ and 4^th^ recipient with and without JUN induction. Scale bar = 500 µm. n=2.

### JUN upregulates CD47 in mouse dermal fibroblasts

Having established and characterized our *Jun*-driven mouse model, we turned to immune therapy. While *Jun* induced CD47 expression in all fibroblast subpopulations, it increased PDL1 expression only in CD26+ and DP fibroblasts in vivo (Fig. 4 A, B, C, S7 A, B). We then explored if immune therapy could eliminate primary mouse dermal fibroblasts in an adaptive transfer model. For this purpose, we treated immunocompromised mice with either a CD47 or a PDL1 inhibitor after transplanting primary mouse dermal fibroblasts under their kidney capsule. Both inhibitors eliminated the graft (Fig. 4 D, E, F, S7 C, D). Regarding CD47, we observed in vitro that JUN decreases and CD47 inhibition increases phagocytosis (Fig. 4 G, H, S8). In regard of PD1/PDL1, we determined PD1 expression on mouse macrophages suggesting a mechanism through which PDL1 blockade is also effective in a T cell-deficient environment such as the immunocompromised NSG mouse (Fig. S7 E, F). Investigating the contribution of macrophages to skin fibrosis initiation, macrophage depletion through a CSFR1 inhibitor ameliorated skin fibrosis (Fig. 4 I, J, K).

**Figure 4.**
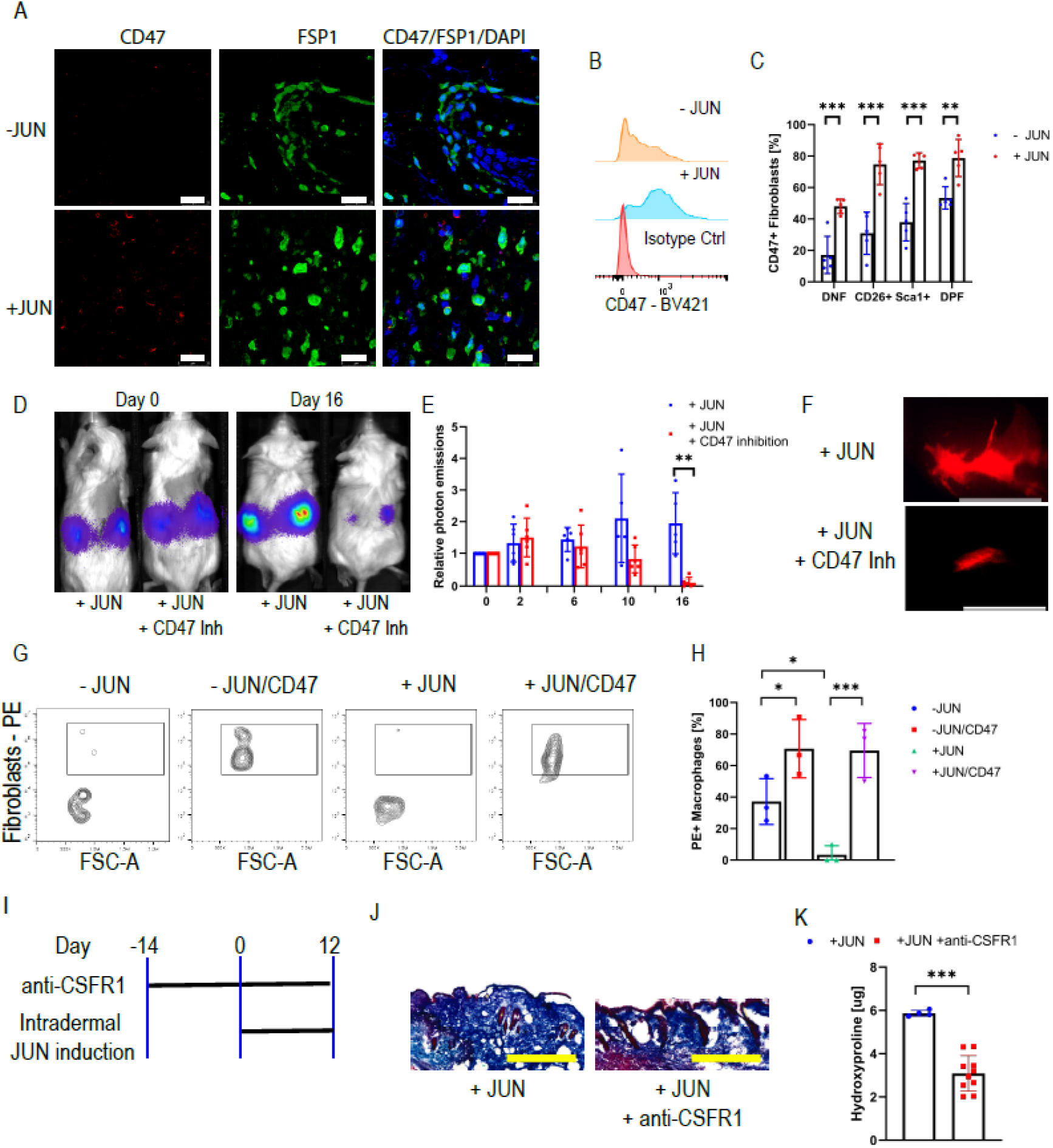
CD47 inhibition eliminates dermal fibroblasts in vivo and in vitro. **(A)** Immunofluorescence stains against CD47 and FSP1 with and without JUN induction. Scale bar = 25 µm. n=5. **(B)** Histogram of CD47 expression in fibroblasts with and without JUN induction. n=5. **(C)** Percentage of CD47 positivity in different fibroblast populations with and without JUN induction. Fisher’s multiple comparions test. *** p < 0.01. n=5. Bar graphs represent means with standard deviations. **(D)** Representative optical images of ectopically transplanted JUN inducible mouse dermal fibroblasts +/- CD47 inhibition. n=4. **(E)** Corresponding quantification of photon emissions. Values are normalized to day 0. Fisher’s multiple comparions test. ** p < 0.01. n=4. Bar graphs represent means with standard deviations. **(F)** Fluorescent graft visualization under the dissection microscope after seven days of CD47 inhibition. Scale bar = 5 mm. n=2. **(G)** Facs Plot for PE/RFP+ CD11b+ macrophages +/- JUN induction +/- CD47 inhibition in an in vitro phagocytosis assay. n=3. **(H)** Corresponding quantification of RFP+ macrophages. Turkey’s multiple comparisons test. * p < 0.05 *** p < 0.01. n=3. Bar graphs represent means with standard deviations. **(I)** Schema of a macrophage depletion trial with subsequent skin fibrosis induction. n=5. **(J)** Corresponding trichrome stains. Scale bar = 500 µm. n=5. **(K)** Corresponding hydroxyproline assay. Two-sided t-test. *** p < 0.001. n=5. Bar graphs represent means with standard deviations.

### Combining CD47 and IL6 inhibition prevents loss in subcutaneous fat tissue

We then investigated if immune therapy targeting either the immune checkpoint PD1/PDL1 or the “don’t-eat-me-signal” CD47 also allows prevention of fibrosis in our *Jun*-induced mouse. As our results had shown that both hedgehog signaling and IL6 contribute to scleroderma, we combined both immune therapies either with vismodegib (CD47/PDL1 inhibition + vismodegib) or with the IL6 inhibitor tocilizumab (CD47/IL6 inhibition). For this purpose, we concomitantly administered these treatment schedules and induced J*un* intradermally for two weeks (Fig. S9 A). Compared to the control group, neither treatments reduced the thickness of the skin or its fibrotic area (Fig. S9 B, C, D). However, the percentage of dermal fat tissue and the fatty area overall was increased with CD47/IL6 inhibition (Fig. S9 E, F). Additionally, both treatment groups demonstrated a decreased cellular dermal infiltrate When examining the cellular infiltrate more closely, both treatments reduced the number of CD3+ cells in the dermis (Fig. S9 G, H). Additionally, both treatments decreased the dermal number of KI67+ cells and prevented the agglomeration of macrophages (Fig. S9 G, H, I, J).

### Therapeutic CD47/IL6 inhibition reverses skin fibrosis

Having evaluated immune therapies in disease initiation, we then tested whether CD47 inhibition reverses skin fibrosis in mice. To study this, we induced skin fibrosis over two weeks before starting treatment with combined CD47/IL6 blockade or IL6 blockade alone (Fig. 5 A, B). Encouragingly, two weeks after treatment initiation, the skin hydroxyproline content in the treated mice was lower than in the control groups (Fig. 5 B, C). Additionally, the fat area was increased under CD47/IL6 inhibition, reversing the skin to an almost normal state (Fig. 5 B, D). Otherwise, the overall infiltrate with immune cells did show no differences between the individual treatment groups (Fig. S10 A - D). Looking for side effects, we additionally harvested other organs. Only the bone marrow demonstrated anemic changes, while the other organs all appeared normal (Fig. S10 E).

**Figure 5.**
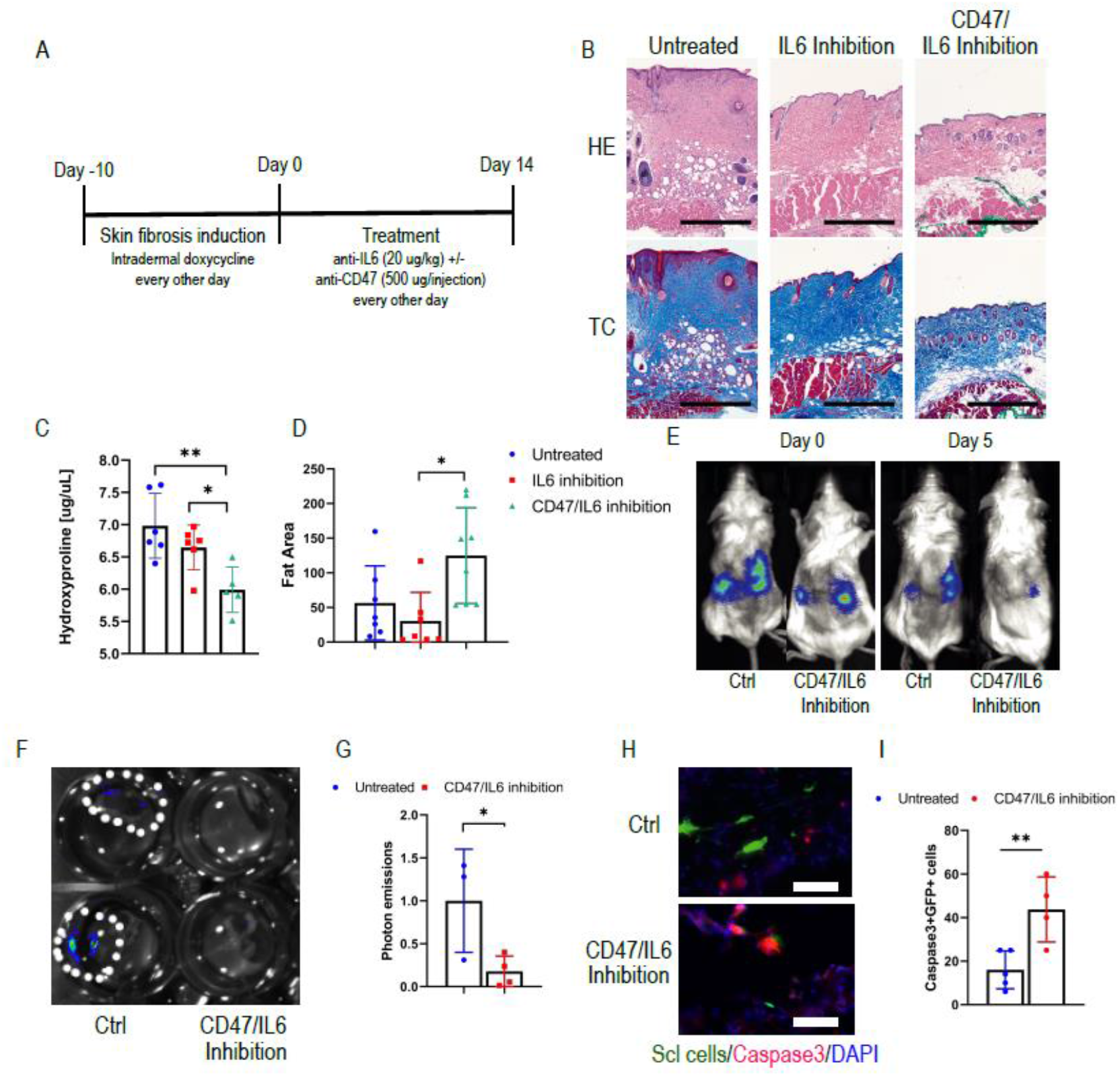
Combined CD47/IL6 inhibition reverses skin fibrosis. **(A)** Schematic outline of the therapeutic trial **(B)** Representative H&E and Trichrome stains of the different groups. Scale bar = 500 µm. n=4. **(C)** Hydroxyproline content of the skin. Turkey’s multiple comparisons test. * p < 0.05 ** p < 0.01. n=6. Graph bars represent means with standard deviations. **(D)** Amount of fat tissue. Values indicate µm^2^/µm skin width. Turkey’s multiple comparisons test. * p < 0.05. n=8. Graph bars represent means with standard deviations. **(E)** Representative optical images of ectopically transplanted GFP/Luciferase-labeled human scleroderma fibroblasts +/- CD47/IL6 inhibition. **(F)** Optical imaging of explanted kidneys on day 5 **(G)** Quantification of photon emissions of explanted kidney grafts normalized to the values of the untreated mice. Two-sided t-test. * p < 0.05. n=3-4. Bar graphs represent means with standard deviations. **(H)** Corresponding Caspase3 staining of kidney grafts. Scale bar = 25 µm. **(I)** Corresponding percentage of Caspase3+ GFP+ fibroblasts. Two-sided t-test. ** p < 0.01. n=4-5. Bar graphs represent means with standard deviations.

### CD47/IL6 inhibition eliminates scleroderma fibroblasts in an adaptive-transfer model

Finally, we evaluated if CD47/IL6 inhibition accelerates the elimination of transplanted human scleroderma fibroblasts in an adaptive transfer model. Two days after transplantation, we started the treatment with CD47/IL6 inhibition and tracked the GFP/Luciferase-labeled cells through optical imaging (Fig. 5 E). Five days after treatment start. Performing optical imaging on the explanted kidneys confirmed decreased optical signals in the treatment group (Fig. 5 F, G). In accordance with the accelerated elimination of the grafts, we additionally observed significantly more apoptotic GFP+ fibroblasts under CD47/IL6 (Fig. 5 H, I). In summary, we were able to demonstrate that blocking CD47 and IL6 prevented the loss in fat tissue during disease initiation, reversed fibrotic skin changes thereafter, and eliminated human scleroderma fibroblasts in an adaptive transfer model.

## Discussion

In this study, we evaluated the efficiency of the “don’t-eat-me-signal” CD47 either alone or in combination with an IL6 inhibitor in a scleroderma mouse model. Systemic sclerosis and its skin manifestation scleroderma are autoimmune diseases and current treatments mainly aim at stopping the progression of the diseases [1]. A true cure with a significant reduction of fibrosis can rarely be achieved. Therefore, a therapy that allows to remove abnormal fibrosis and fibroblasts would represent a breakthrough.

We split this study into three different parts with three main questions, whereby the first two parts cleared the way for the third part in which we addressed the efficiency of CD47 inhibition for fibrosis. The first part served to determine whether *JUN* contributes to human scleroderma and in the second part, we addressed mechanisms that drive skin fibrosis in our *Jun*-driven mouse model.

Regarding the role of *JUN* in human scleroderma, we demonstrate that scleroderma fibroblasts activate *JUN* on the protein level. In accordance with that, the promotor accessibility of *JUN* is increased in primary human scleroderma fibroblasts compared to normal skin fibroblasts. We additionally demonstrate that the promotors of the hedgehog-associated genes *GLI1* and *PTCH1* are more readily accessible in scleroderma fibroblasts. *JUN* knockout reduced the promotor accessibility of *PTCH1* and *GLI1*, and vice versa, the hedgehog inhibitor vismodegib reduced the promotor accessibility of *JUN*. This points towards the interconnection between *JUN* and hedgehog signaling. Importantly, the promotors of CD47 and PDL1 were more easily accessible in scleroderma, suggesting that the abnormal fibroblasts use “don’t-eat-me-signals” and immune checkpoints as a protective mechanism against the host’s immune system. Additionally, the interleukin 6 was more readily accessible, a finding that corresponds with a recent study in which IL6 activated profibrotic pathways in explanted scleroderma samples [24]. At the end of this part, we demonstrate that stiffer conditions on a hydrogel induce JUN activation, both in scleroderma and pulmonary fibroblasts. Though this does not answer which role *JUN* plays during disease initiation, it points – in conjunction with the results from our *Jun*-driven mouse model - towards the detrimental role *JUN* in disease progression.

In the second part of this study, we aimed at characterizing our mouse model in order to better interpret our results from the treatment with the CD47 inhibitor as we had to identify whether our *Jun*-driven mouse model overlaps with human scleroderma. Intradermal *Jun* induction caused dermal fibrosis and a loss in subcutaneous fat tissue. Additionally, it reduced CD31+ endothelial cells and increased the infiltration with CD3+ cells, hence exhibiting fibrosis, vasculopathy and inflammation – three of four hallmarks of scleroderma. Except the decrease in CD31+ cells, *Jun* induction mimicked the changes after bleomycin injection. Interestingly, both *Jun* induction and bleomycin led to the same extent of JUN activation. In accordance with our results from the human fibroblasts on the hydrogel, a possibility is that bleomycin first leads to a temporary fibrosis and stiffening of the skin that then leads to JUN activation. According to the *Jun*-driven mouse model JUN activation alone is enough to cause dermal fibrosis. Once activated JUN could then maintain and worsen skin fibrosis through IL6/pStat3 signaling. This mechanism could also apply to human disease though we could not finally address this question in this study. As similar roles have been demonstrated for other AP1 family members such as *Fra2* and *JunB*, the deregulated activation of only one of the various AP1 family members may be sufficient to promote skin fibrosis [25].

In the following experiments, we show that JUN expands CD26+ fibroblasts and decreases DP fibroblasts in the beginning. When induced over a longer time course, CD26+ fibroblasts then start to decrease and DN fibroblasts expand significantly. These results suggest that during fibrosis initiate CD26+ fibroblasts first increase and then finally differentiate into CD26-fibroblasts., In the following experiments, we demonstrate that CD26+ fibroblast activate hedgehog signaling and that *Jun* induction even further increases this activation. Importantly, inhibiting the hedgehog pathway with vismodegib reverses two previous findings – both the expansion of CD26+ fibroblasts and the dermal fibrosis. In conclusion, these data demonstrate that JUN leads to dermal fibrosis through the distinct activation of hedgehog signaling in CD26+ fibroblasts.

There is ongoing debate about the cell from which fibrosis stems [26, 27]. In an adaptive transfer model, we show that *Jun* expands dermal fibroblasts and supports graft survival independent of their dermal niche. Additionally, *Jun* induction allowed to detect transplanted fibroblasts up to the fourth recipient in a serial transplantation model, strongly indicating that JUN mediates increased self-renewal to fibroblasts. These results support the hypothesis that a clonal expansion of fibroblasts might be the source of fibrotic changes in scleroderma.

In the main part of study, we assessed if immune therapy, targeting either CD47 or PDL1, either in combination with vismodegib or in combination with IL6 blockade allows to prevent or reverse skin fibrosis. *Jun* induction increased CD47 in all and PDL1 in two of the fibroblast populations. Blocking either CD47 or PDL1 in the adaptive transfer model led to the elimination of the graft while the graft continued to grow without immune therapy. In accordance with its role as a “don’t-eat-me-signal” CD47 inhibition increased phagocytosis in vitro. Regarding the PD1/PDL1 axis, we demonstrated in accordance with recent publications that mouse macrophages in our models express PD1 as well, suggesting that PDL1-inhibition in the T-cell deficient NSG mouse works through its interaction with PD1 on macrophages. There is ongoing debate over the role of macrophages in fibrosis initiation. In this study, we observed that the depletion of macrophages through a CSFR1 inhibitor ameliorated skin fibrosis.

We then determined the efficiency of CD47 inhibition in the syngeneic mouse. We first demonstrated that CD47/IL6 inhibition did not decrease the content of connective tissue in the skin in a prospective study. However, the loss of fat tissue with stiffening of the skin is a hallmark of scleroderma and CD47/IL6 increased the dermal fat content. Importantly, we then moved to a therapeutic study in which we first induced *Jun* over two weeks before starting treatment. To determine to which extent IL6 blockade contributes to changes in skin fibrosis in our mouse model, we added a IL6 blockade only group. Strikingly, CD47/IL6 significantly decreased skin fibrosis and increased dermal fat, in comparison both to the untreated and the IL6 blockade only group. Exempt anemic changes, we did not observe any histological side effects under CD47/IL6 inhibition. In accordance with these findings, CD47/IL6 blockade accelerated the elimination of human scleroderma fibroblasts and increased apoptosis in an adaptive transfer model.

Our macrophage depletion study and our therapeutic study point demonstrate that macrophages can act as a double-edged sword. On one hand they worsen fibrosis initiation, on the other hand they can remove fibrotic tissue. To protect themselves from phagocytosis, fibroblasts upregulate CD47 during disease initiation and progression, thereby enhancing their survival. Through CD47 blockade, fibroblasts become more accessible to macrophages, and can finally be eliminated through phagocytosis.

Clinically, immune therapies can lead to immune-related adverse events and safety of immune checkpoint inhibition in general, and in autoimmune disease in particular, is a primordial question [28, 29]. Although data are still incomplete, retrospective studies suggest that these patients can be treated safely and effectively with immune therapies [29]. Both CD47 and IL6 inhibition are normally well tolerated. CD47 inhibition can cause anemia and infections are more common in patients treated with the IL6 antibody tocilizumab [30, 31]. However, these adverse events are rarely high-grade. In accordance with that, exempt anemic changes in the bone marrow, CD47/IL6 treated mice did not show any other histologic side effects under CD47 and IL6 inhibition.

Despite its thorough evaluation of CD47 inhibition, open questions remain. One of these questions concerns the pathogenesis of scleroderma. Mouse models can only partly mimic diseases and are unsuitable to reflect the complete complexity of human disease [32]. This is especially true for complex diseases such as scleroderma whose pathogenesis is incompletely understood. There is good evidence from the literature and this study that JUN contributes to scleroderma and fibrotic diseases in general. All the same, other factors will contribute as well. A *Jun*-driven scleroderma mouse model is genetically biased and will miss contributing factors to scleroderma. Therefore, our results concerning the pathogenesis of scleroderma need to be read with caution.

Despite these limitations, our study is a proof of concept that combined immunotherapy can reverse fibrotic skin conditions in mice. The pathogenesis leading to fibrosis in different conditions may vary, however, they share an end stage with abnormal and persistent fibroblasts leading to significant morbidity and even death. Current therapies mainly stop disease progression but do not reverse fibrosis. Combining CD47 inhibition to enhance phagocytosis and IL6 blockade to suppress underlying detrimental inflammatory processes, in contrast, could lead to a true healing of fibrosis. Regarding the good safety profiles of currently available CD47 and IL6 blocking agents, we believe that our study gives enough evidence to safely try combined immunotherapy with CD47 and IL6 inhibition in patients with highly fibrotic and stable, non-progressive scleroderma.

## Materials and Methods

### Animal Studies

Animal trials were approved by the Stanford Administrative Panel on Laboratory Animal Care (#30911) and mice were kept in the facilities of the Stanford Veterinary Service Center on a regular diet and experiments were performed according to the approved protocol.

### Husbandry

JUN mice were kept in the facilities of the Veterinary Service Center at Stanford University and had a B6/129 background. Nod.Scid.Gamma mice were purchased from the Jackson Laboratory. Mice were kept on a standard diet. Female and male mice were used. When different sexes were used for individual experiments, groups were sex-matched. Mice were not backcrossed and between 6 and 12 weeks of age during experiments.

### Genotyping

To determine the genotype, we harvested tissue from the tail of newborn mice on day 10. We digested the DNA with Quickextract (Lucigen Corporation) at 68°C for 90 minutes, followed by heat inactivation at 98° C for five minutes. We ran the genotyping PCR for the Rosa26 and the collagen status with the Phusion® High Fidelity DNA Polymerase (New England Biolabs) and the same primers as described previously and indicated in the Supplementary Table 2 [6].

### Adaptive transfer under the kidney capsule

After anesthetizing mice, we areas over both flanks were shaved. After creating a flank cut the subcutaneous tissue was bluntly removed from the underlying soft tissue. The abdominal wall was incised and the kidney luxated out of the abdominal cavity. After piercing the kidney and detaching the renal capsule from the renal tissue, 50,000 to 200,000 cells suspended in matrigel were injected under the kidney capsule. Afterwards, the kidney was replaced into the abdominal cavity and the abdominal wall and the skin were separately sutured.

### Administration of vismodegib, PDL1 inhibitor, CD47 antibody and CSFR1 antibody

We administered vismodegib (30 mg/kg bodyweight) (Selleckchem) intraperitoneally two times daily. The CD47 antibody (Bio X Cell) was given every other day. The first dosage was 100 μg, followed by 500 μg. For PDL1 inhibition, we injected 100 μl (2.5 mg/ml) of HAC anti-PD1 daily [33]. To deplete macrophages, we injected 400 µg of the CSFR1 antibody (Bio X Cell) every other.

### Intradermal and systemic JUN induction

To induce JUN locally, we injected 20 μl of doxycycline (MilliporeSigma) (2 mg/ml) intradermally on the back. For systemic JUN induction, we injected doxycycline intraperitoneally (20 μg/g body weight). In both systems, we performed the injections every other day.

### Luciferase-based optimal imaging

We intraperitoneally injected 100 цl of luciferin substrate (15 mg/ml) (Biosynth). 15 minutes later, we performed optical imaging with the Lago optical imaging system (Spectral imaging instruments) and analyzed the images with the Aura Software from the same manufacturer.

### Harvesting of mouse skin for subsequent cell cultures or flow cytometry

Mice were euthanized with CO_2_. Their backs were shaved, washed and disinfected with 70% EtOH. After excising the back skin, tissue was minced with scissors, followed by a digestion step in DMEM + 10 % PS, supplemented with 40 μl/ml of Liberase (Roche), for 30 minutes at 37° in the cell incubator on a shaker. The digestion reaction was quenched with DMEM + 10 % FCS. The medium and tissue specimens were filtered through a 70 μm cell strainer (Falcon). Cells were then either used for flow cytometry or cell culture.

### Isolation and maintenance of primary mouse dermal fibroblast cultures

After filtering cells and tissue as described previously, cells were washed two times with PBS. The supernatant was removed and cell pellets and tissue parts were transferred into a culture dish. Cells were kept in DMEM + 5 % HPL supplemented with Ciprofloxacin over five days. Medium was changed on the first and third day. When being 50 % confluent, cells were spliited.

### Isolation of primary human scleroderma and pulmonary fibroblast cultures

Human fibroblasts were obtained discarded fresh lungs tissues from de-identified patients. The tissue was minced and filtered through 70 μm filters, centrifuged at 600 g for 5 min to remove non-homogenized pieces of tissue. Tissue homogenate was treated with ACK lysing buffer (Thermo Fisher) for 10-15 min, centrifuged for 600 g, washed twice in DMEM with 10% fetal bovine serum (Gibco) and plated at a density of approximately 500,000 cells/cm2 in DMEM with 10% fetal bovine serum, 1% penicillin/streptomycin (Thermo Fisher Scientific) and Ciprofloxacin 10 μg/mL, Corning) and kept in an incubator at 37° C 95% O2 / 5% CO2. Media was changed after 24h and cells were cultured until 80-90% confluent before each passage.

### Cell culture maintenance

All cell cultures were kept in DMEM supplemented with HPL. Cells were regularly checked for signs for infection and splitted once being more than 80% confluent. For splitting, cell cultures were washed with PBS and incubated with Trypsin (Gibco) for 5 minutes. The reaction was then quenched with DMEM + HPL. Cell suspensions were spun down and reapplied on cell dishes.

### Phagocytosis assay

Peritoneal macrophages were harvested from non-JUN inducible B6 mice. After euthanizing the mice, the skin above the abdomen was cut to expose the peritoneum. The abdominal cavity was then flushed with 5 ml of cold 50 mM EDTA without disrupting vessels. The injected fluid was then aspirated. After adding 10 ml of PBS, the cell suspension was centrifuged for 5 minutes at 300 g, followed by another washing step with PBS and a centrifugation step. The harvested cells were then plated into a 10 ml dish in regular medium. After two hours, the medium was exchanged and M-CSF (20 µg/ml) (MilliporeSigma) was added. After two days, the medium was exchanged with fresh M-CSF. In the meantime, JUN inducible fibroblasts were prepared and labeled with a RFP plasmid. On day 2, JUN was induced in one group by adding doxycycline (1 µg/ml) to the cell culture medium. On day 3, macrophages and fibroblasts with and without JUN induction were harvested and counted. The fibroblast populations were then split again, one group with CD47 inhibition and one group without CD47 inhibition. Cells with CD47 inhibition were then incubated for one hour with the CD47 antibody (Bio X cell) on ice in FACS buffer while the cells were kept on ice in FACS buffer as well. For measuring phagocytosis through flow cytometry, 25,000 macrophages and fibroblasts were mixed in individual wells of a 96-well plate. After a two hour incubation period on a shaker in a regular cell incubator, wells were washed with cold PBS, followed by trypsinization for 10 minutes. FACS buffer added, the plate was centrifuged, followed by another washing step with FACS buffer. After spinning down the plate, cells were incubated with a CD11b and CD45 antibody (BioLegend) for 45 minutes on ice. Afterward, cells were washed and resuspended in FACS buffer, followed by flow analysis in a CytoFlex Flow cytometer (Beckman Coulter). For analysis, CD45+CD11b+ cells were gated and the percentage of RFP/PE+ cells determined. For immunofluorescence, macrophages were plated on fibronectin (MilliporeSigma) coated glass slides (VWR). After 45 minutes, fibroblasts were added and incubated with macrophages for one hour. After that, slides were vigorously washed three times with cold PBS, followed by a regular stain of cells plated on glass slides as described elsewhere in the method section.

### Hydrogel preparation

The wells of a 24-well glass bottom plates (Mattek) were incubated with 2 M NaOH for one hour. After a washing step with ddH_2_O, the wells were incubated with 500 µl of 2% Aminopropyltriethoxysilane (MilliporeSigma) (diluted in 95% EtOH). The wells were rinsed with ddH_2_O, followed by incubating the wells with 500 µl of 0.5% Glutaraldehyd (MilliporeSigma) for 30 minutes. The wells were then rinsed with ddH_2_O and dried at 60° C. For a stiffness of 70 kPa, 46.25 µl of 40% Acrylamid (MilliporeSigma) were mixed with 55.5 µl of 2% bis-acrylamid (MilliporeSigma) and 83.25 µl of ddH_2_O. Then, 1.2 µl of 10% Ammonium persulfate (MilliporeSigma) and 0.8 µl of TEMED (MilliporeSigma) were added to the mixture before. Each well of the plate was coated with 4 µl of the mixture and a coverslip (Glaswarenfabrik Karl Hecht GmbH). After 20 minutes, the coverslip was removed with tweezers. 500 µl of 50 mM HEPES (MilliporeSigma) were added to the wells. After sterilizing the plate for 1 hour under UV light, 200 µl of 0.5% SANPAH crosslinker (ProteoChem) were added to each well. After activating the crosslinker on UV light for ten minutes, wells were washed with 50 mM HEPES two times. A 1/15 solution of Matrigel (MilliporeSigma) diluted in 50 mM HEPES were added to the plate. The plate was incubated overnight at room temperature. Before being used, the plate was rinsed with 50 mM HEPES and incubated with regular cell culture medium for 30 minutes at 37° C in the incubator.

### Hydroxyproline assay

We determined the hydroxyproline content with a Hydroxyproline assay kit (MilliporeSigma) according to the manufacturer’s specifications. 10mg of tissue was minced with scissors and homogenized in 100µL of deionized water. 100µL of 12M HCL was added to each sample and incubated at 120 °C for 3 hours. Samples were centrifuged at 10,000 g for 3 minutes and 25 µL of the supernatant was plated into a 96-well and subsequently incubated at 60 °C until all liquid was evaporated. Standards and reagents were prepared as instructed in the manufacturer’s protocol. 100 µL of the chloramine T oxidation buffer mixture was added to each sample and standard and incubated for 5 minutes at room temperature. 100 µL of diluted DMAB reagent was added to each sample and standard and incubated in a 60 °C water bath for 90 minutes. Absorbance was read at 560 nm. All samples and standards were performed in technical duplicates.

### Preparation of human platelet lysate (HPL) and HPL-containing medium

Expired human platelets were obtained from the Stanford Blood Bank. Then, platelets were lysed through five quick freeze-thaw cycles. Platelet lysates were then spun down at 4,000 g for 10 minutes, aliquoted into 15 ml tubes and stored at −80° C. For preparing cell mediums, platelet lysates were warmed up, then spun down at 4,000 g for 10 minutes, and sequentially filtered through 0.80, 0.45 and 0.22 μm filters. The final medium, containing DMEM, 5 % HPL, 1 % Penicillin/Streptomycin and 2 Units of Heparin/ml, was then filtered through a 0.22 μm filter and stored at 4° C.

### Flow cytometry and sorting

For live cells, we washed single cell suspensions with FACS buffer (PBS + 2 % FBS + 1 % Penicillin/Streptomycin + 1 mM EDTA + 25 mM HEPES) and then stained the cells with the primary antibodies for 45 minutes. Cells were washed and subsequently resuspended with FACS buffer. For intracellular stainings, cells were fixed with BD Wash/Perm (Becton, Dickinson and Company), followed by the same steps used for the live cells, with the exception of using BD Wash/Perm (Becton Dickinson and Company) instead of the FACS buffer. For flow cytometry, we either used the CytoFlex Flow Cytometer (Beckman Coulter) for analysis only, or the BD FACS Aria III (Becton, Dickinson and Company) for analysis and sorting. We performed the data analysis with the newest version of FlowJo (FlowJo, LLC).

### Tissue fixation and hematoxylin staining

We kept harvested tissue in 10% formalin overnight. Tissue was then submitted to the Stanford Human Pathology/Histology Service Center for paraffin-embedding and cutting. We deparaffinized and rehydrated the tissue slides with xylene and a descending ethanol row. After washing the slides in PBS, we incubated them in hematoxylin (American MasterTech) for 4 minutes, then in bluing reagent (ThermoFisher Scientific) for 2 minutes and in Harleco® (MilliporeSigma) for 2 minutes. Slides were dehydrated with ethanol and xylene (MilliporeSigma) and mounted wit Permount® (ThermoFisher Scientific).

### Trichrome Staining

We used a One Step Trichrome Stain Kit (American MasterTech). After deparaffinization and rehydration, the tissue was incubated in Bouin’s Fluid overnight, followed by Modified Mayer’s Hematoxylin for seven minutes and One Step Trichrome Stain for five minutes. Slides were dehydrated with ethanol and xylene and covered with Permount® (ThermoFisher Scientific).

### Immunofluorescence Staining of paraffin-embedded sections

When using paraffin-embedded tissue, we first deparaffinized and rehydrated the tissue. Then, we performed antigen retrieval with a citric acid buffer in a pressure cooker for 15 minutes, followed by blocking with 5 % serum. Sections were incubated with the primary antibody over night at 4° C. After washing in PBST, we incubated the sections with the secondary antibody at room temperature for 30 minutes under agitation. Sections were washed, counterstained with DAPI and mounted with fluoromount (SouthernBiotech). Images of histological slides were obtained on a Leica Eclipse E400 microscope (Leica) equipped with a SPOT RT color digital camera model 2.1.1 (Diagnostic Instruments).

### Immunofluorescence Staining of OCT-embedded sections

For OCT sections, harvested tissue was fixed in 4% PFA for at least two hours, followed by an incubation in 30% sucrose overnight. Tissue was then embedded in OCT and stored at −80° C. Thereafter, the OCT block was cut into 10 µm sections. Sections were stored at -20° C. Before staining, sections were dried for at least 15 minutes at room temperature. Sections were rehydrated in PBS. For intranuclear stains, sections were incubated for 10 minutes in TritonX at room temperature. Otherwise, cells were blocked with 5% for one hour. The remaining steps were the same as for the immunofluorescence stains of paraffine-embedded tissue.

### Immunofluorescence Staining of cells plated on glass slides

For plating cells on glass slides (VWR), areas were encircled on glass slides with a fat pen (Vector Laboratories), followed by sterilization for at least one hour under UV light. Then, fibronectin (20 µg/ml PBS) was added. After incubating the slides for one hour in a cell incubator at 37° C, the fibronectin solution was aspirated and the slides were dried in the cell culture cabinet. Cells were added onto the glass slides, and incubated for different amount of times in regular cell culture medium. Thereafter, the glass slides were washed in PBS, followed by fixing the cells with 4% PFA at room temperature for ten minutes. The slides were washed again in PBS, followed by permeabilization in TritonX for ten minutes for intranuclear stainings. Otherwise, cells were blocked with 5% serum for one hour. Afterwards, the staining procedure equaled the staining protocol of the paraffin-embedded slides.

### Immunohistochemistry

When using paraffin-embedded tissue, we first deparaffinized and rehydrated the tissue. Then, we performed antigen retrieval with a citric acid buffer in a pressure cooker for 15 minutes, followed by blocking with 5 % serum. Sections were incubated with the primary antibody over night at 4° C. After washing in PBST, we incubated the sections with the secondary HRP-antibody at room temperature for one 30 minutes under agitation. Sections were washed in PBST and ddH_2_O followed by incubation for 15-20 minutes in AEC Peroxidase Substrate (Vector Laboratories). Sections were washed in ddH_2_O and incubated in Modified Mayer’s Hematoxylin for 4 Minutes. After washing in ddH_2_O, slides were mounted with fluoromount (SouthernBiotech).

### RNA extraction, cDNA and quantitative polymerase chain reaction

FACS purified cells were sorted into trizol (ThermoFisher Scientific). For RNA extraction, we added chloroform and centrifuged the tubes. We transferred the upper phase to a new tube and added 70 % EtOH. Afterwards, the complete volume was added to the columns of the RNEasy MiniElute Cleanup Kit (Qiagen). Between the next three centrifugation steps, we washed the columns with 80% EtOH (+H_2_O), 80% EtOH (+RPE) and 70%EtOH. After letting the membranes slightly dry, we added water onto the membranes, centrifuged the columns and measured the RNA quantity and quality of the flow through with a NanoDrop 2000 (ThermoFisher Scientific). For cDNA creation, we used the iScript™ Advanced cDNA Synthesis Kit (Bio-Rad) according to the manufacturer’s specifications. Up to 1 μg of RNA was transformed into cDNA. We then ran qPCR on a 7900 HT Fast-Time PCR System (Applied Biosystems). Reactions contained 2.5 µl of H_2_O, 5 µl of PowerUp^TM^ SYBR^TM^ Green Master Mix (Applied Biosystems), 1 µM of Primers (0.5 µl) and 2 µl of cDNA.

### Western Blotting

Cells in a well of a 6-well-plate were lysed with 150–200 μl of urea buffer. Cell lysates were sonicated and centrifuged. Protein concentrations were determined with the Pierce BSA Protein Assay (ThermoFisher Scientific). 10 μg of protein were mixed with loading dye and heated up to 99° C for five minutes. Protein lysates were added to a Bolt 4-12% Bis-Tris-Plus Gel (Invitrogen). After 30 minutes, proteins were transferred to a nitrocellulose membrane for 90 minutes. Membranes were blocked with 5% milk powder in PBST for 30 minutes, followed by incubating the membranes with the primary antibody diluted in the blocking buffer for 1 hour at room temperature. Membranes were washed three times with PBST for five minutes each. Membranes were incubated with the secondary antibody diluted in the blocking buffer at room temperature, followed by three washing steps in PBST. Membranes were incubated with Luminata Forte Western HRP Substrate (MilliporeSigma) for five minutes and developed on UltraCruz® Autoradiography films (Santa Cruz Biotechnology). Membranes were stripped with Restore^TM^ Western Blot Stripping Buffer (ThermoFisher Scientific) for five minutes. Membranes were then blocked, incubated with primary and secondary antibodies and developed as described in this section.

### Primary antibodies

Flow cytometry: **CD3** (Biolegend, #100209, Clone 17A2), **CD4** (Biolegend, #100422, Clone GK1.5), **CD11b** (BD, #554411, Clone M1/70), **CD11c** (Biolegend, #117324, Clone N418), **CD25** (Biolegend #102035, Clone PC61), **CD26** (Biolegend, #137805, Clone H194-112), **CD31** (BD, #553373, Clone MEC 13.3), **CD45** (Biolegend, #103110, Clone 30-F11), **CD47** (Biolegend, #127527, Clone Miap301), **CD326** (Biolegend, #118218, Clone G8.8), F4/80 (Biolegend, #123116, Clone BM8), **PDL1** (Biolegend, #124312, Clone 10F.9G2), **Sca1** (Biolegend, #108114), **phospho c-Jun** (Ser73) (CST, #32705, Clone D47G9) Immunohistochemistry/Immunofluorescence: **Adiponectin** (abcam, #ab22554), **CD3** (abcam, #ab5690), **CD11b** (Novus, #NB110-89474), **CD26** (abcam, #ab28340), **CD26** (R&D, #AF954), **CD31** (Dako, #m0823), **CD47** (FisherScientific, #14-0479-82, Clone B6H12), **CD47** (R&D, #AF1866), **CD68** (Agilent, #GA60691-2, Clone KP1), **Cleaved Caspase 3** (CST, #96645, Clone 5A1E), **FSP1** (MilliporeSigma, #07-2274), **FSP1** (Abcam, #ab58597), **Collagen 1** (Abcam, #ab34710), FSP1 (MilliporeSigma, #07-2274), **Ki67** (abcam, #ab15580), **PD1** (Cell marque, #315M-96, Clone NAT105), **PD1** (R&D, #AF1021), **PDL1** (R&D, #AF1019), **phospho c-Jun** (Ser73) (CST, #32705, Clone D47G9).

Western **c-Jun** (CST, #9165S, Clone 60A8), **FSP1** (MilliporeSigma, #07-2274), GAPD (GeneTex, #627408, Clone GT239), **phospho c-Jun** (Ser73) (CST, #32705, Clone D47G9), **phospho Stat3** (CST, #9131S).

### Secondary antobodies

**AF488 Goat anti-rabbit** (CST, #44125), **AF594 Goat anti-rabbit** (Invitrogen, #A-11012), **AF594 Donkey anti-goat** (Novus, #NBP1-75607), **HRP Goat anti-rabbit** (abcam, #205718).

### ATAC-Seq Library preparation, sequencing, and data preprocessing

The ATAC-seq was performed as discribed before. Briefly, 50000 cells were collected and washed with cold PBS and lysed using 0.1% NP40 in resuspension buffer. Tn5 transposition of nuclei pellets was carried out at 37°C for 30 min, using the DNA sample preparation kit from Nextera (Illumina). The reaction was purified using QIAGEN MinElute columns and then amplified for 8-15 cycles to produce libraries for sequencing. ATAC-seq libraries were sequenced on Illumina HiSeq 4000. ATAC-seq pair-end reads were trimmed for Illumina adaptor sequences and transposase sequences using Kundaje ATAC_pipelines. The libraries were initially sequenced on a Miseq sequencer and analyzed using a custom script to determine the enrichment score, only libraries that had the highest score above the threshold (>5) were chosen for deeper sequencing. Two independent, biological replicates were sequenced per sample. The data have been uploaded to the Gene Expression Omnibus (GEO) under the accession number GSE151943.

### Deep sequencing data analysis

Differentially accessible peaks from the merged union peak list were selected with the DESeq2 package from bioconductor using raw read counts of each samples using log2 fold change > 1, and p value < 0.05. The read counts of the differential peaks in each sample were further normalized by Z-score transformation. Peak genomic annotation was performed by HOMER package. Hierarchical clustering was used to cluster the peaks and samples. The results was presented as heatmap by Morpheus from Broad Institute.

### Statistical analysis

We used the newest version of Graphpad prism (Graphpad Software Inc) for creating graphs and running statistical analyses. When two values were compared, a two-sided student’s t-test was used, if more than two values were analyzed. When more than two values were directly compared to each other, we used the Turkey’s multiple comparisons test. When two values were compared over different time points, without comparing the time points to each other, we used Fisher’s multiple comparisons test. Generally, experiments included at least three independent values from two independent experiments. P values below 0.05 were considered as statistically significant. Regarding naturally occurring higher variation in animal trials, we determined before the experiment to exclude the highest and lowest values (when n was at least 8) or the two highest and two lowest values (when n of 16 was reached). All primary data will be made available upon request.

## Author contributions

TL: Study conception, study interpretation, running and analyzing all experiments except ATAC Seq studies and Western Blots, writing manuscript LC: Running and analyzing ATAC Seq studies MK: Participation in running and analyzing animal trials shown in figures 6 – 8, writing manuscript TC: Running all Western Blots CM: Writing manuscript TS: Writing manuscript GW: Study conception, data interpretation, final manuscript approval.

## Acknowledgments

We thank Pauline Chen and Shirley Kwok for their help and elaborate expertise in embedding and cutting various forms of tissue and Norman Cyr for his help in taking histological pictures.

## Supplementary Materials

**Supplementary Figure 1.**
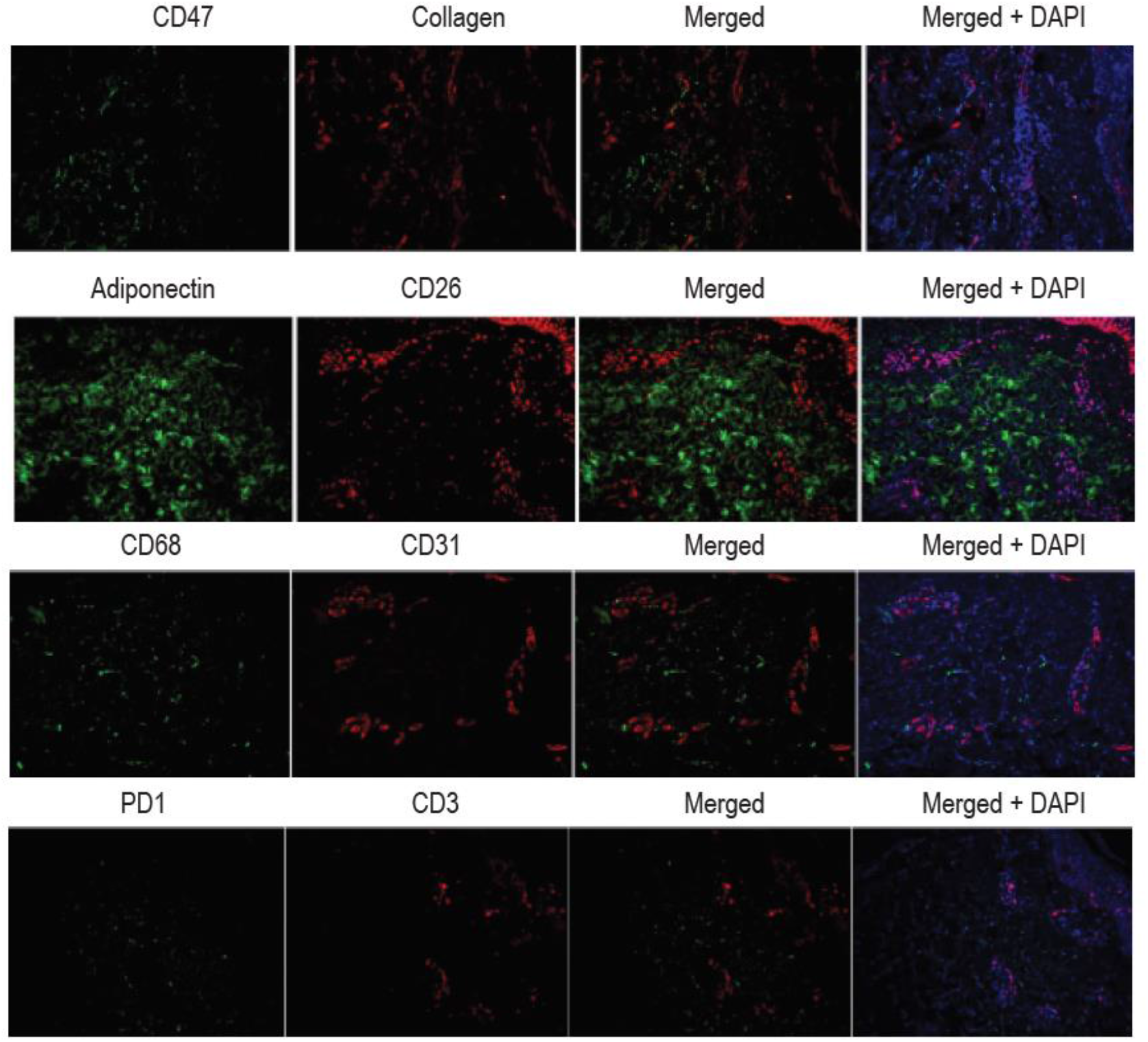
Expression of different markers in human scleroderma samples. Chromatin accessibility changes in human scleroderma samples. Representative images of human scleroderma skin sections stained against different markers.

**Supplementary Figure 2.**
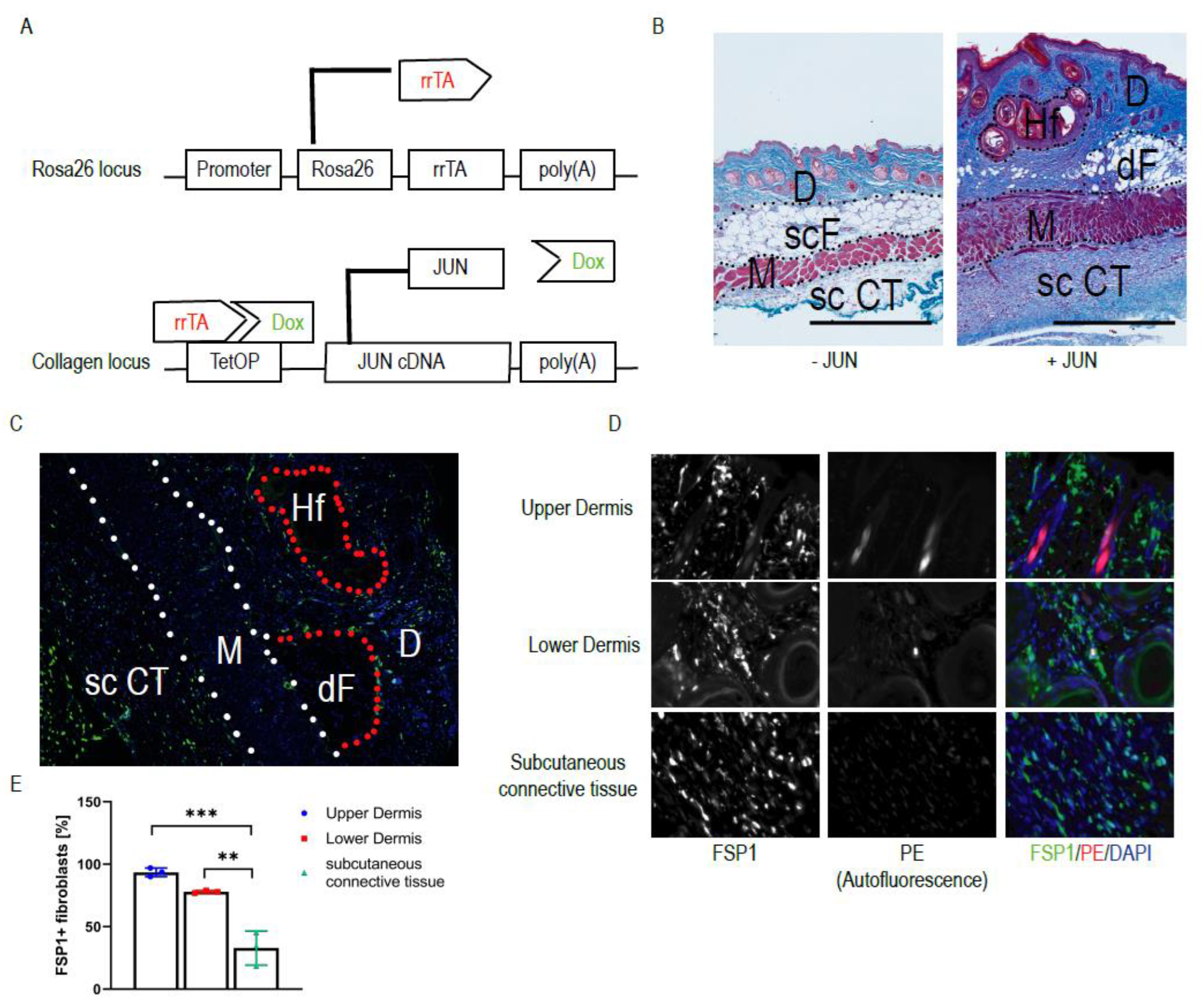
Jun expands FSP1+ fibroblasts throughout the skin and the underlying tissue. **(A)** Genetic modifications of the JUN-inducible mouse. rrTA = reverse tetracycline transactivator, TetOP = Tetracycline/doxycycline-responsive operator **(B)** Representative Trichrome-stained whole skin sections without (-JUN) and with JUN induction (+JUN). D = Dermis, scF = subcutaneous fat, M = subcutaneous muscle, sc CT = subcutaneous connective tissue, dF = dermal fat, Hf = Hairfollicle. Scale bar = 500 µm. **(C)** Whole skin section after Jun induction (corresponding to the section B). Green = FSP1+, blue = DAPI. D = Dermis, M = subcutaneous muscle, sc CT = subcutaneous connective tissue, dF = dermal fat, Hf = Hairfollicle **(D)** Quantification of FSP1+ fibroblasts in the upper dermis, the lower dermis and the subcutaneous connective tissue. Indicated are the percentages of FSP1+ cells among all spindle-shaped fibroblasts. Turkey’s multiple comparisons test. ** p < 0.01 *** p < 0.001. Scale = 500 µm. n=3. Bar graphs represent means with standard deviations. **(E)** Representative stains against FSP1 in the upper and lower dermis and the subcutaneous connective tissue after Jun induction.

**Supplementary Figure 3.**
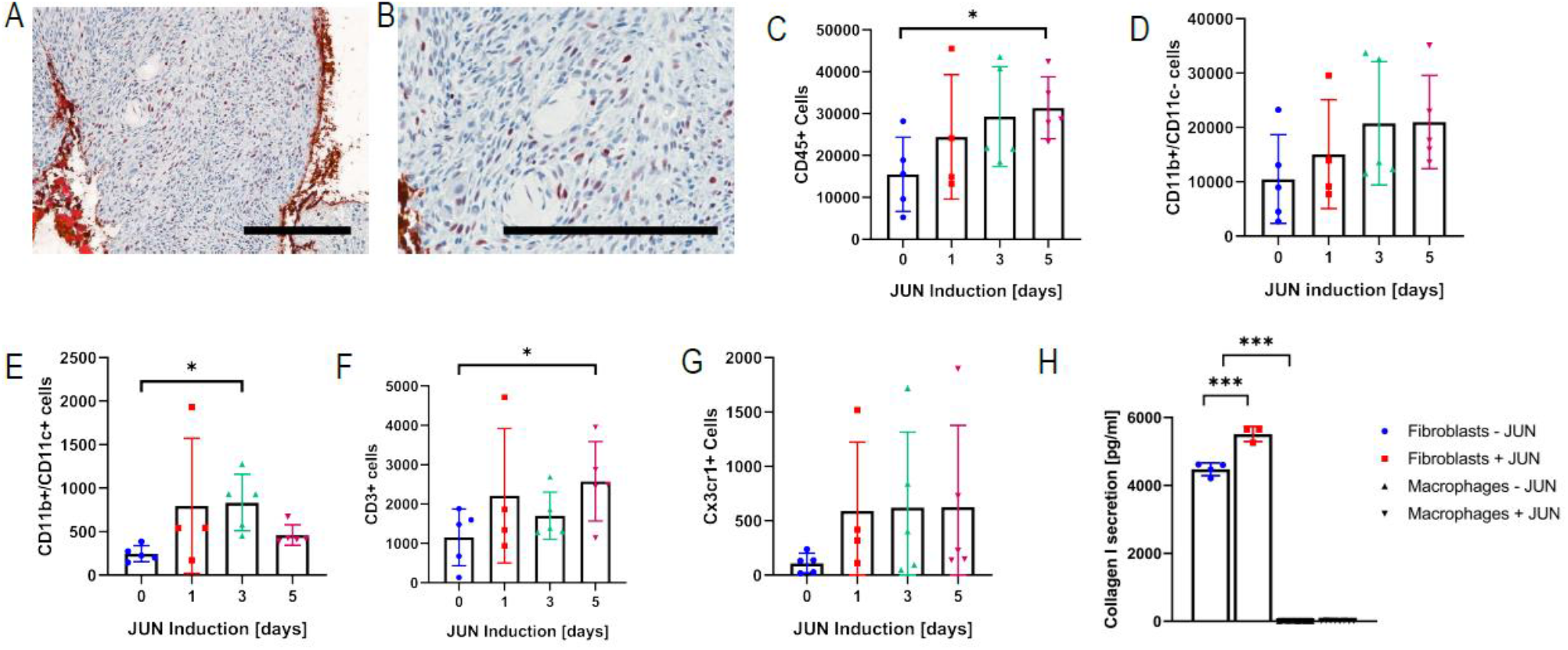
Skin immune filtration under JUN induction. **(A)** Representative IHC stain against pJUN. Scale bar = 200 µm. **(B)** Representative IHC stain against pJUN. Scale bar = 200 µm. **(C)** Quantification of CD45+ cells under JUN induction over up to 5 days. Turkey’s multiple comparisons test. * p < 0.05. n=4-5. Bar graphs represent means with standard deviations. **(D)** Quantification of myeloid CD11b+ cells under JUN induction over up to 5 days. Fisher’s multiple comparisons test. n=4-5. Bar graphs represent means with standard deviations. **(E)** Quantification of dendritic CD11b+CD11c+ cells under JUN induction over up to 5 days. Turkey’s multiple comparions test. * p < 0.05. n=4-5. Bar graphs represent means with standard deviations. **(F)** Quantification of CD3+ T cells under JUN induction over up to 5 days. Turkey’s multiple comparisons test. * p < 0.05. n=4-5. Graph bars represent means with standard deviations. **(G)** Quantification of hematopoietic Cx3cr1+ cells under JUN induction over up to 5 days. Fisher’s multiple comparisons test. n=4-5. Graph bars represent means with standard deviations. **(H)** In vitro collagen 1 secretion of fibroblasts and macrophages +/- JUN induction. One-way ANVOA * p < 0.05 *** p < 0.001 (n=4). Bar graphs represent means with standard deviations.

**Supplementary Figure 4.**
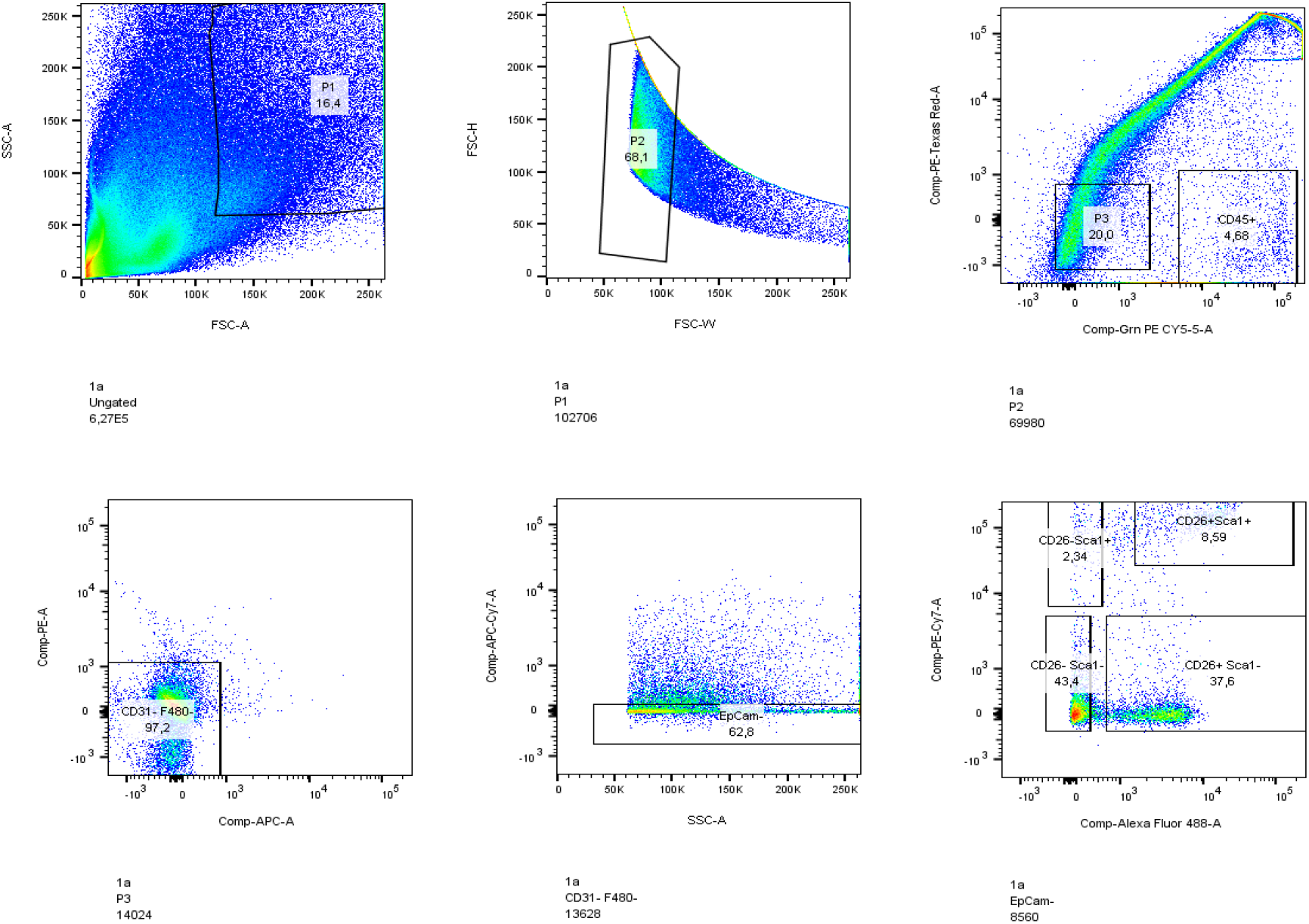
Fibroblast gating strategy. After identifying cells and then single cells, hematopoietic (CD45+) and dead cells (PI) are excluded. In a next step, macrophages (F4/80+) and endothelial (CD31+) cells are excluded. After removing epithelial (CD326+) cells, fibroblasts are divided, based on their expression of CD26 and Sca1.

**Supplementary Figure 5.**
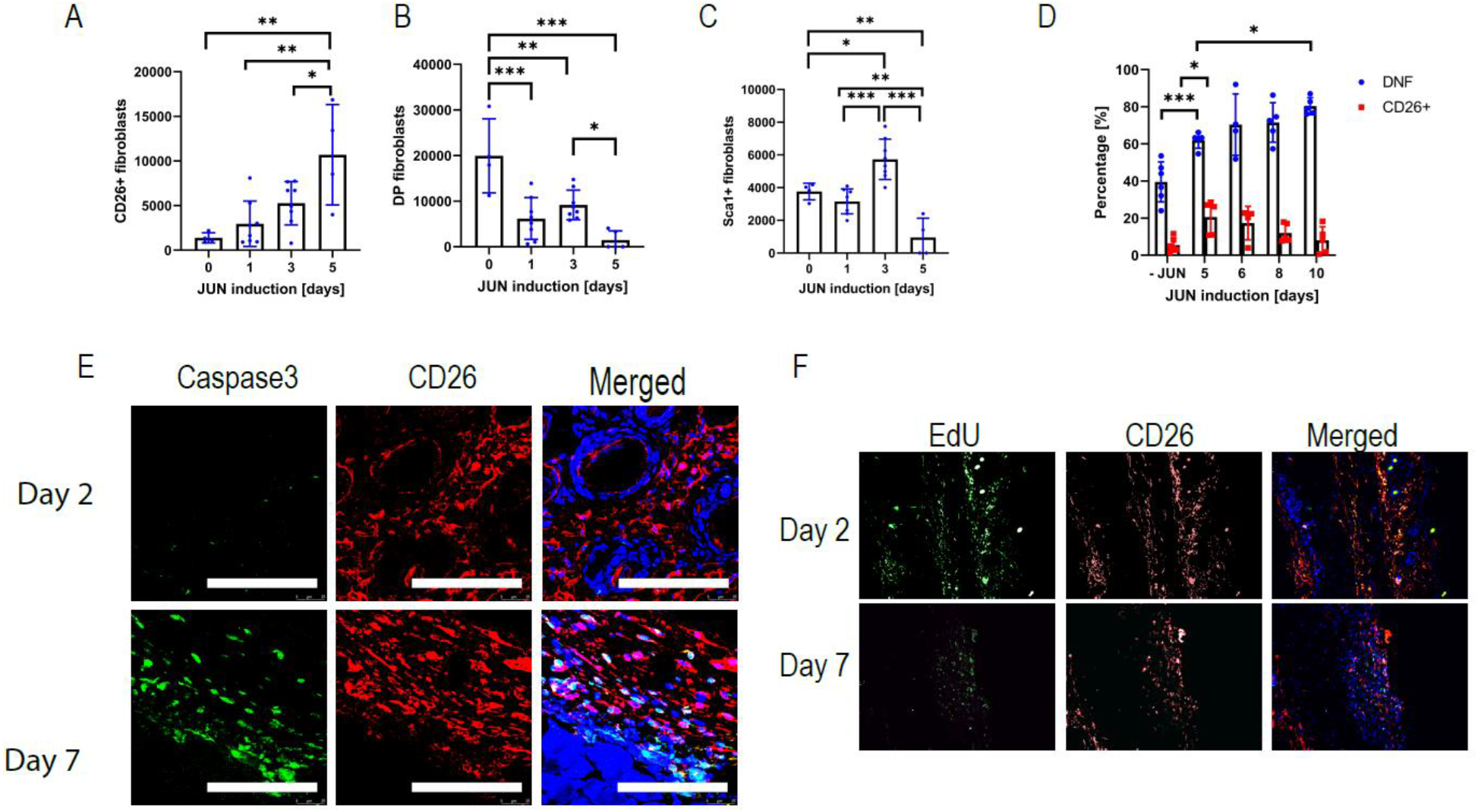
JUN initially expands CD26+ fibroblasts. **(A)** Quantification of CD26+ fibroblasts, over five days of JUN induction. Turkey’s multiple comparisons test. * p < 0.05 ** p < 0.01. n=4-5. Bar graphs represent means with standard deviations. **(B)** Quantification of DPF over five days of JUN induction. Turkey’s multiple comparisons test. *** p < 0.001. n=5. Bar graphs represent means with standard deviations. **(C)** Quantification of Sca1+ fibroblasts over five days of JUN induction. Turkey’s multiple comparisons test. ** p < 0.01 *** p < 0.001. n=5. Bar graphs represent means with standard deviations. **(D)** Quantification of CD26+ fibroblasts and DP fibroblasts (DPF) over up to 10 days of JUN induction. Fisher’s multiple comparisons test. * p < 0.05 ** p < 0.01 *** p < 0.001. n=4-6. Bar graphs represent means with standard deviations. **(E)** Representative immunofluorescence stains against Caspase3 and CD26 two and seven days after JUN induction. Scale bar = 100 µm. **(F)** Representative immunofluorescence stain against CD26 and corresponding EdU visualization two and seven days after JUN induction.

**Supplementary Figure 6.**
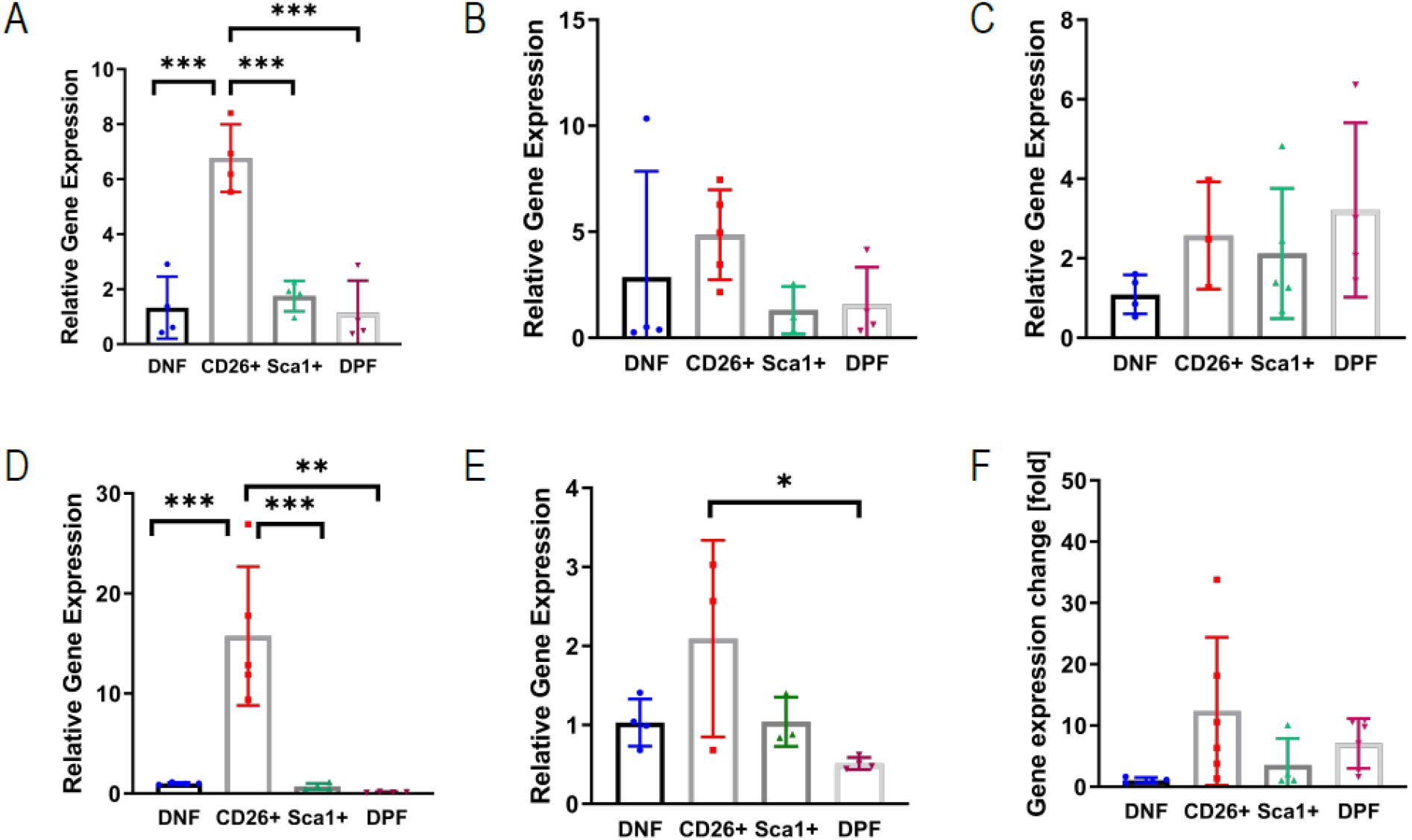
CD26+ fibroblasts activate hedgehog signaling. Normalized qPCR data from facs purified fibroblast populations from non JUN-induced JUN mice. The values for each gene are compared to the mean value of the DN fibroblasts. (A) Gli1. Turkey’s multiple comparisons test. *** p < 0.001. n =4. Bar graphs represent means with standard deviations. (B) Gli2. Turkey’s multiple comparisons test. n=3-5. Bar graphs represent means with standard deviations. (C) Gli3. Turkey’s multiple comparisons test. n=3-5. Bar graphs represent means with standard deviations. (D) Ptch1. Turkey’s multiple comparisons test. ** p < 0.01 *** p <0.001. n=3-5. Bar graphs represent means with standard deviations. (E) Kif7. Turkey’s multiple comparisons test. * p < 0.05. n=3-4. Bar graphs represent means with standard deviations. (F) Smo. Turkey’s multiple comparisons test. n=4-6. Bar graphs represent means with standard deviations.

**Supplementary Figure 7.**
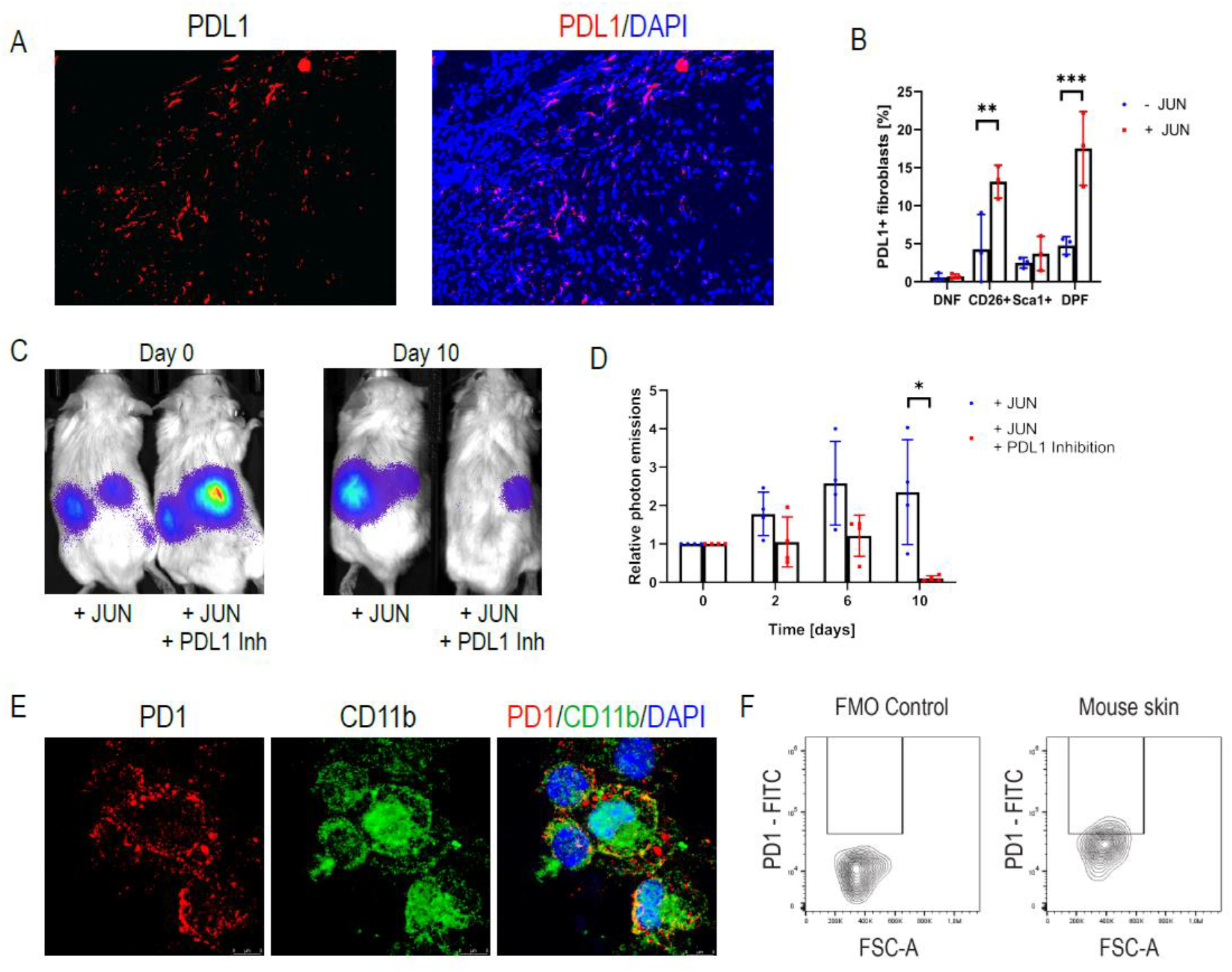
PDL1 inhibition eliminates ectopic fibroblasts. **(A)** Immunofluorescence stains against PDL1 after local JUN induction in skin after 14 days **(B)** PDL1 expression in different subsets of fibroblasts with and without JUN induction. Turkey’s multiple comparisons test. ** p < 0.01 *** p < 0.001. n=3. Bar graphs represent means with standard deviations. **(C)** Representative optical images of ectopically transplanted JUN inducible fibroblasts +/- PDL1 inhibition. n=4. **(D)** Corresponding quantification of photon emissions. Turkey’s multiple comparisons test. * p < 0.05. n=4. Bar graphs represent means with standard deviations. **(E)** Immunofluorescence stains against PD1 and CD11b on macrophages harvested from the peritoneum. **(F)** FACS plots of PD1 expression in CD45+CD11b+ blood cells. n=2. Graph bars represent means with standard deviations.

**Supplementary Figure 8.**
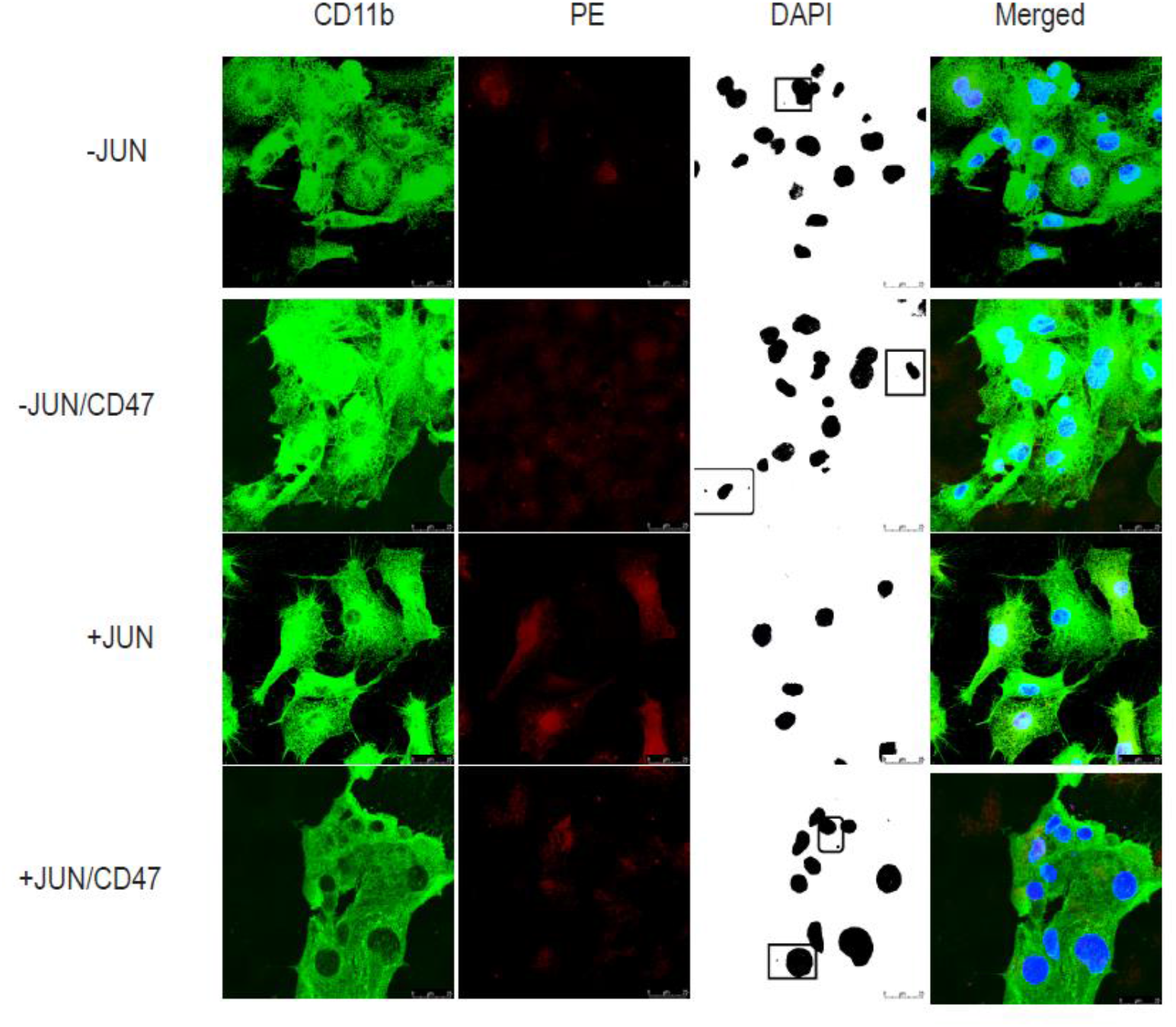
CD47 inhibition increases phagocytosis of dermal fibroblasts in vitro. Images are taken with a confocal microscope. RFP+ target cells are detected in the PE channel. Macrophages who have fully digested target cells are marked by small isolated DNA pieces. Boxes in the DAPI represent macrophages with additional DNA pieces as signs of advanced phagocytosis.

**Supplementary Figure 9.**
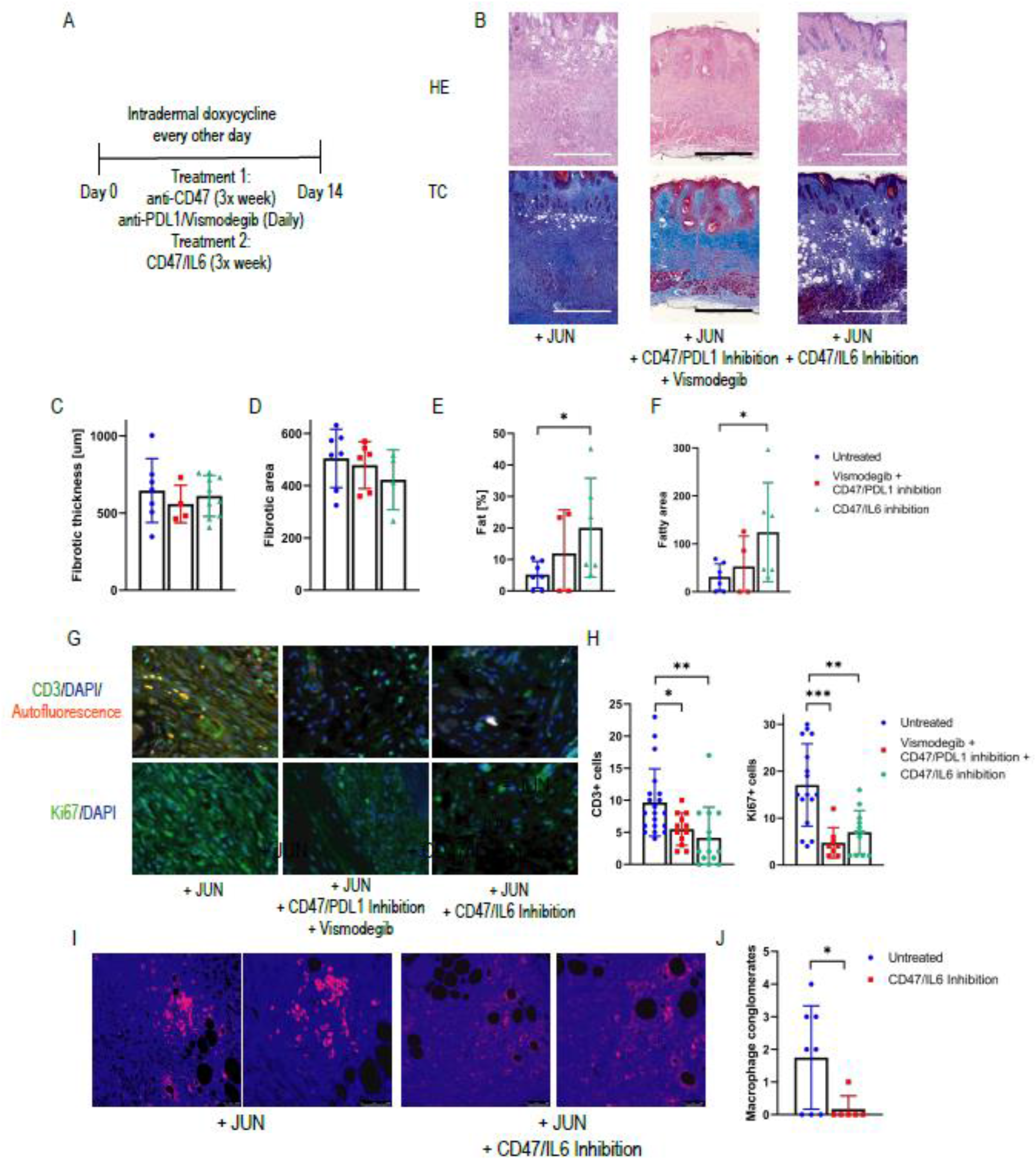
Combining CD47 and IL6 inhibition prevents loss in subcutaneous fat tissue. **(A)** Experimental outline **(B)** Representative H&E and Trichrome skin stains of untreated mice, mice under CD47/PDL1 inhibition and vismodegib, and mice under CD47/IL6 inhibition. Scale = 500 µm. Bar graphs represent means with standard deviations. **(C)** Thickness of the dermal fibrotic/connective tissue in µm. Fisher’s multiple comparisons test. n=4-7. Bar graphs represent means with standard deviations. **(D)** Area of the fibrotic tissue in untreated and treated samples, values indicate µm^2^/µm skin width. Fisher’s multiple comparisons test. n=4-7. Bar graphs represent means with standard deviations. **(E)** Percentage of dermal fat, compared to the overall dermal area, in treated and untreated samples. Turkey’s multiple comparisons test. * p < 0.05. n=4-7. Bar graphs represent means with standard deviations. **(F)** Area of dermal fat tissue in untreated and treated samples, values indicate µm^2^/µm skin width. Turkey’s multiple comparisons test. * p < 0.05. n=4-7. Bar graphs represent means with standard deviations. **(G)** Representative stains against Ki67 and CD3. Counterstains with DAPI. **(H)** Quantification of CD3+ and Ki67+ stains. Indicated are the number of positive cells/high power view (63x). Turkey’s multiple comparisons test. * p < 0.05 ** p < 0.01 *** p < 0.001. n=8-20. Bar graphs represent means with standard deviations. **(I)** Representative pictures of CD11b+ cells in skin fibrosis +/- CD47/IL6 inhibition. **(J)** Quantification of macrophage agglomerates determined by more than 20 macrophages/High power view in each sections. Two-sided t-test * p < 0.05. n=8. Bar graphs represent means with standard deviations.

**Supplementary Figure 10.**
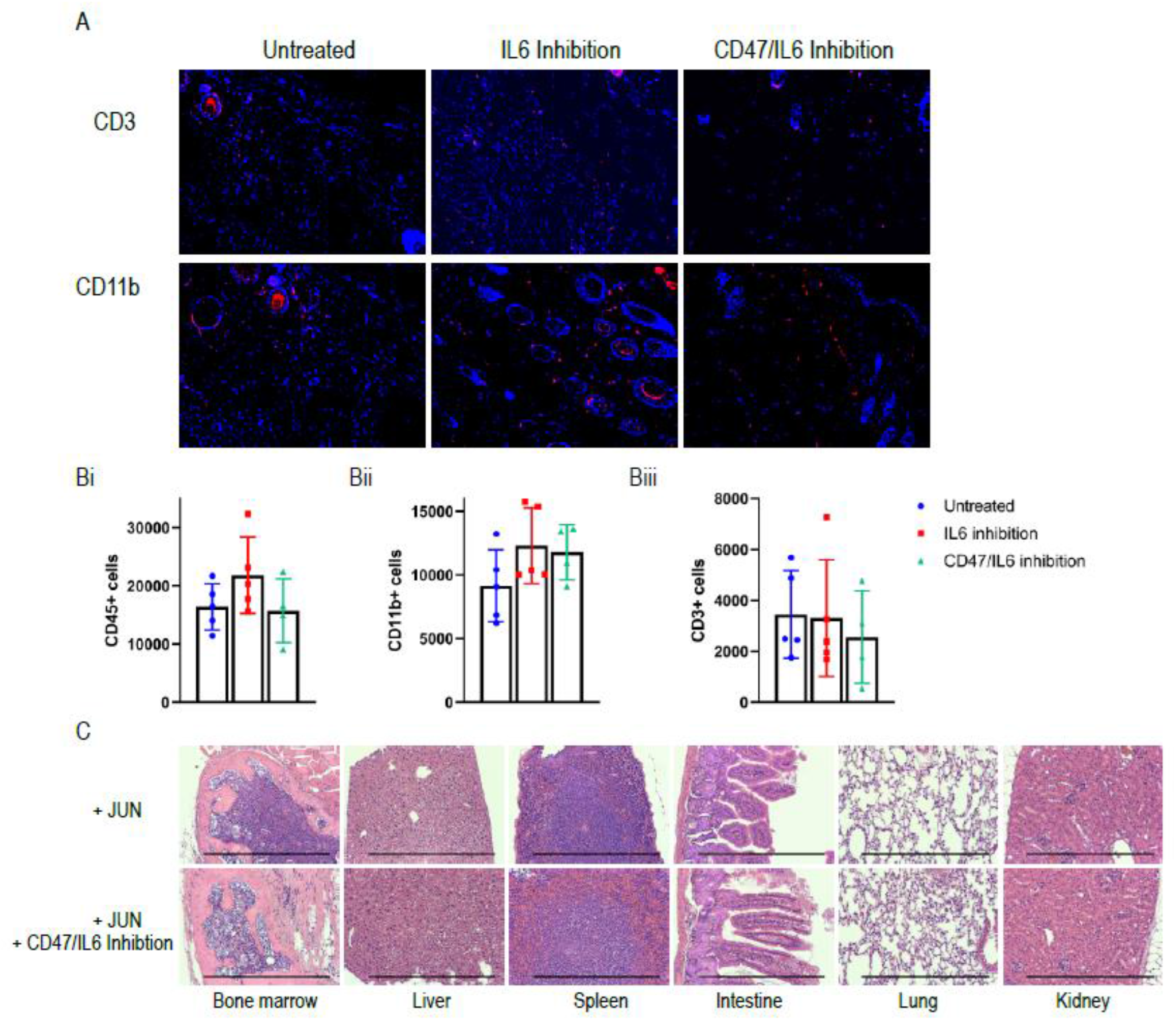
Immune infiltrate in the therapeutic study. **(A)** Representative immunofluorescence stains against CD3 and CD11b in the three groups (Untreated, IL6 inhibition only and CD47/IL6 inhibition) **(B)** Quantification through flow cytometry for CD45+ cells. Turkey’s multiple comparisons test. n=4. Bar graphs represent means with standard deviations. **(C)** Quantification through flow cytometry for myeloid CD11b+ cells. Turkey’s multiple comparisons test. n=4. Bar graphs represent means with standard deviations. **(D)** Quantification for CD3+ T cells. Turkey’s multiple comparisons test. n=4. Bar graphs represent means with standard deviations. **(E)** Corresponding organ sections from the untreated and the CD47/IL6 inhibition group. Flow cytometry numbers represent number of cells/100,000 live cells.

**Supplementary Table 1.**
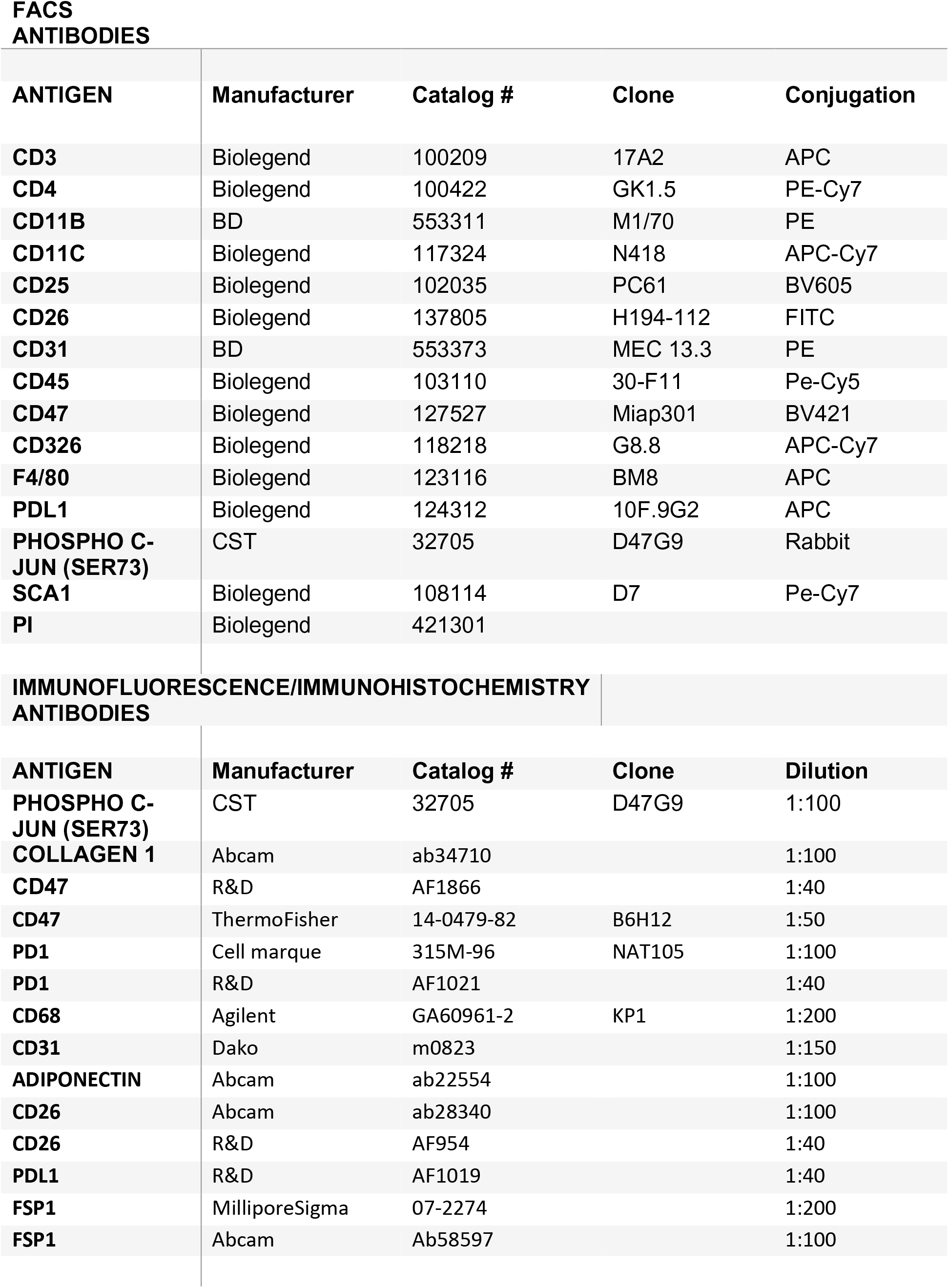
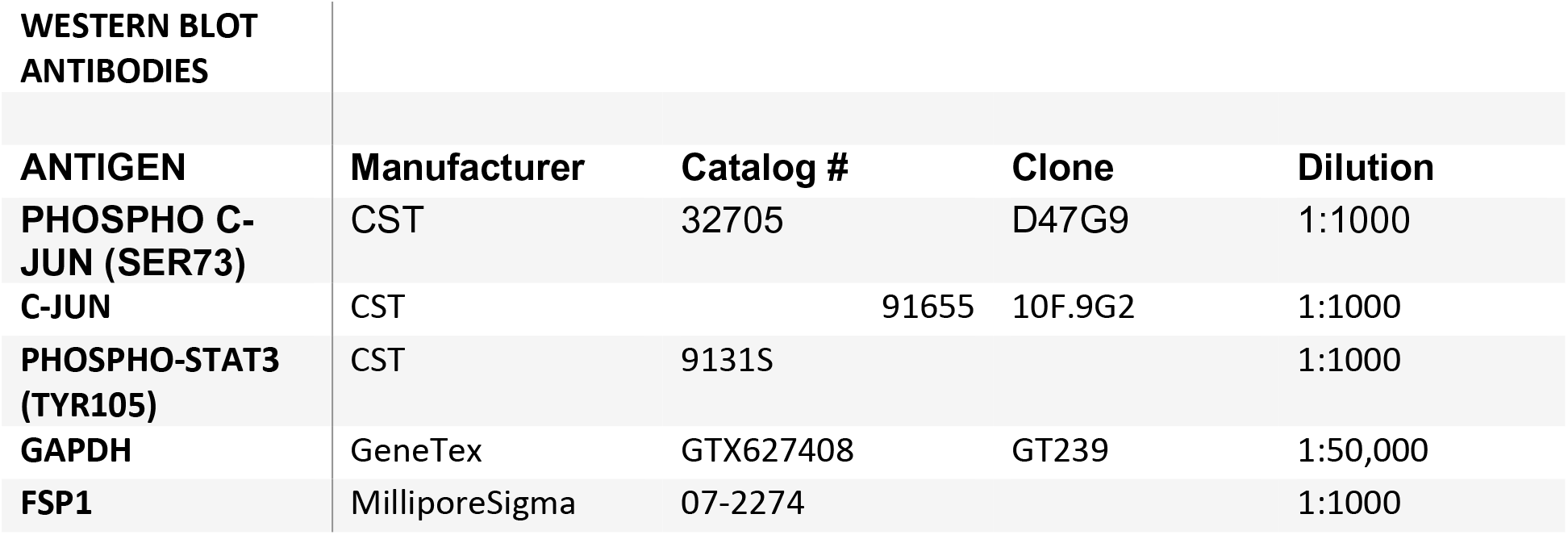
Primary antibodies.

**FACS ANTIBODIES**
| **ANTIGEN** | **Manufacturer** | **Catalog #** | **Clone** | **Conjugation** |
| --- | --- | --- | --- | --- |
| **CD3** | Biolegend | 100209 | 17A2 | APC |
| **CD4** | Biolegend | 100422 | GK1.5 | PE-Cy7 |
| **CD11B** | BD | 553311 | M1/70 | PE |
| **CD11C** | Biolegend | 117324 | N418 | APC-Cy7 |
| **CD25** | Biolegend | 102035 | PC61 | BV605 |
| **CD26** | Biolegend | 137805 | H194-112 | FITC |
| **CD31** | BD | 553373 | MEC 13.3 | PE |
| **CD45** | Biolegend | 103110 | 30-F11 | Pe-Cy5 |
| **CD47** | Biolegend | 127527 | Miap301 | BV421 |
| **CD326** | Biolegend | 118218 | G8.8 | APC-Cy7 |
| **F4/80** | Biolegend | 123116 | BM8 | APC |
| **PDL1** | Biolegend | 124312 | 10F.9G2 | APC |
| **PHOSPHO C-JUN (SER73)** | CST | 32705 | D47G9 | Rabbit |
| **SCA1** | Biolegend | 108114 | D7 | Pe-Cy7 |
| **PI** | Biolegend | 421301 |  |  |

**IMMUNOFLUORESCENCE/IMMUNOHISTOCHEMISTRY ANTIBODIES**
| **ANTIGEN** | **Manufacturer** | **Catalog #** | **Clone** | **Dilution** |
| --- | --- | --- | --- | --- |
| **PHOSPHO C-JUN (SER73)** | CST | 32705 | D47G9 | 1:100 |
| **COLLAGEN 1** | Abcam | ab34710 |  | 1:100 |
| **CD47** | R&D | AF1866 |  | 1:40 |
| **CD47** | ThermoFisher | 14-0479-82 | B6H12 | 1:50 |
| **PD1** | Cell marque | 315M-96 | NAT105 | 1:100 |
| **PD1** | R&D | AF1021 |  | 1:40 |
| **CD68** | Agilent | GA60961-2 | KP1 | 1:200 |
| **CD31** | Dako | m0823 |  | 1:150 |
| **ADIPONECTIN** | Abcam | ab22554 |  | 1:100 |
| **CD26** | Abcam | ab28340 |  | 1:100 |
| **CD26** | R&D | AF954 |  | 1:40 |
| **PDL1** | R&D | AF1019 |  | 1:40 |
| **FSP1** | MilliporeSigma | 07-2274 |  | 1:200 |
| **FSP1** | Abcam | Ab58597 |  | 1:100 |

**Supplementary Table 1. Primary antibodies.**
| WESTERN BLOT ANTIBODIES |  |  |  |  |
| --- | --- | --- | --- | --- |
| ANTIGEN | Manufacturer | Catalog # | Clone | Dilution |
| PHOSPHO C-JUN (SER73) | CST | 32705 | D47G9 | 1:1000 |
| C-JUN | CST | 91655 | 10F.9G2 | 1:1000 |
| PHOSPHO-STAT3 (TYR105) | CST | 9131S |  | 1:1000 |
| GAPDH | GeneTex | GTX627408 | GT239 | 1:50,000 |
| FSP1 | MilliporeSigma | 07-2274 |  | 1:1000 |

**Supplementary Table 2.**
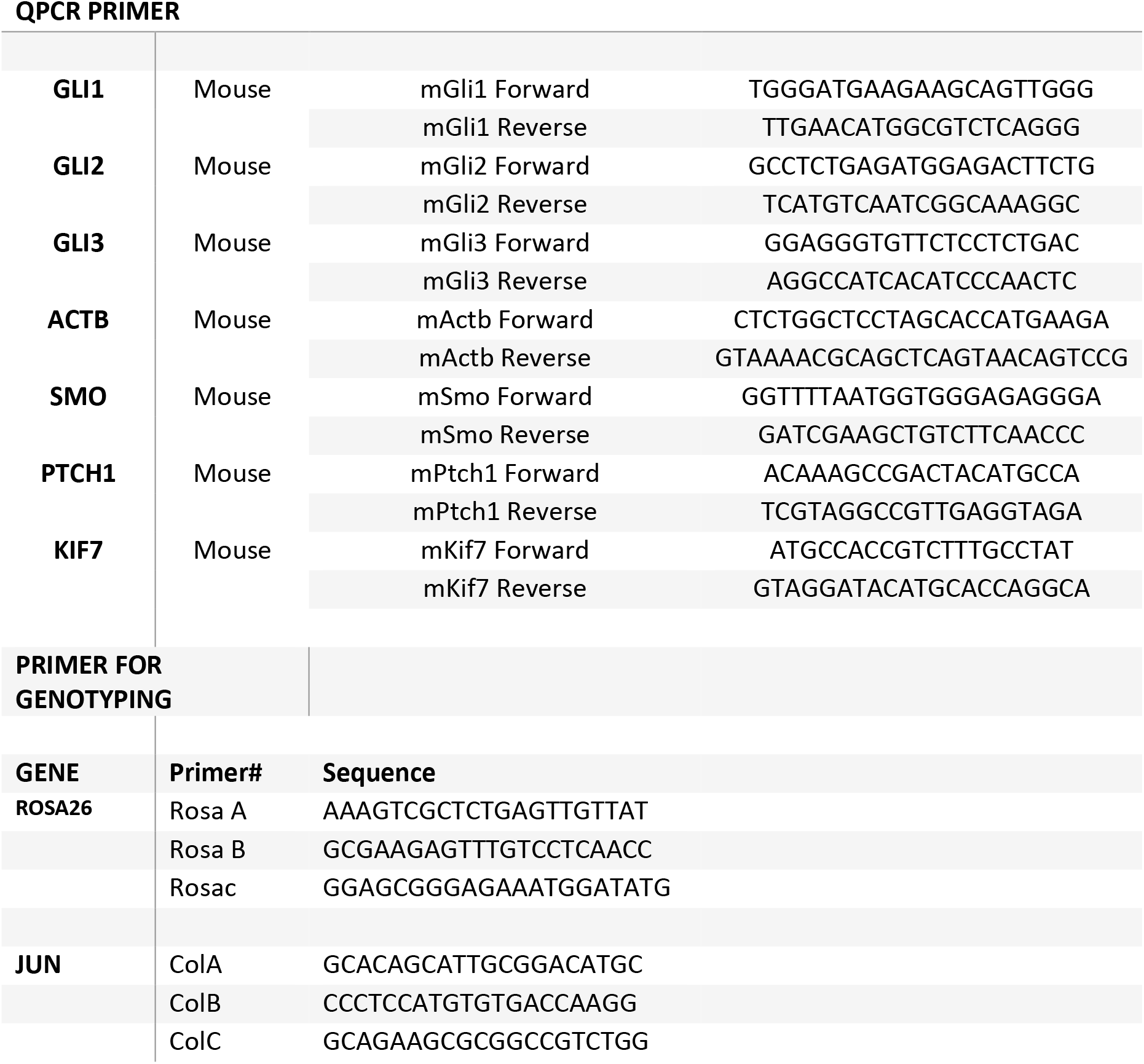
Primer sequences.

